# Microbial induction of MHC-II expression in colon cancer cells overcomes immunotherapy resistance and limits metastasis

**DOI:** 10.64898/2026.06.30.735621

**Authors:** Charlie Chung, Elif Ozcelik, Jialing Zhang, Zeli Shen, Onur Eskiocak, Yihan Qin, Jill Habel, Kadir Ozler, Santhilal Subhash, Zakaria Aminzada, Gagan Dev, Libia Garcia, Scott K. Lyons, Timothy W. Hand, David E. Rivadeneira, James G. Fox, Peter M.K. Westcott, Meri Rogava, Semir Beyaz

## Abstract

Colorectal cancer remains a major cause of cancer mortality, and most microsatellite stable tumors derive little benefit from immune checkpoint blockade. Here, we identify a microbiome-dependent mechanism that converts immune-refractory colorectal cancer into a more immunologically responsive state. Using orthotopic mouse models spanning distinct genetic and immunologic contexts, we show that a Helicobacter-containing microbiome suppresses primary tumor growth and limits metastasis. This protective state is associated with increased intratumoral lymphocyte infiltration and stronger effector programs. Mechanistically, microbial exposure induces MHC class II expression in colon cancer cells to promote anti-tumor immunity. Tumor-intrinsic loss of CIITA abrogates microbial protection, whereas enforced CIITA expression is sufficient to increase intratumoral T cell accumulation, restrict progression and metastasis, and sensitize microsatellite-stable tumors to PD-1 and CTLA-4 blockade. In human microsatellite-stable patient-derived organoids, increased cancer-cell MHC-II enhanced interactions with autologous immune cells and increased tumor cell apoptosis. Together, these findings define a microbiome-cancer cell antigen presentation axis that restrains metastasis and overcomes immunotherapy resistance in colorectal cancer.

## INTRODUCTION

Colorectal cancer (CRC) remains a leading cause of cancer mortality, driven predominantly by metastatic disease that is refractory to standard-of-care therapies (Shiels et al., 2025; Siegel et al., 2026a; Siegel et al., 2026b). Although immune checkpoint blockade has produced durable benefit in mismatch repair-deficient (dMMR) and microsatellite instability-high (MSI-H) CRC, most CRCs are mismatch repair-proficient and microsatellite stable (MSS), and the advanced stages of this disease subset derive little benefit from current PD-1 or CTLA-4 based immunotherapy modalities (Le et al., 2015; Sharma et al., 2023). Clinical efforts to extend checkpoint blockade to MSS metastatic CRC have thus far shown limited efficacy, underscoring the need to identify mechanisms that convert poorly inflamed tumors into lesions capable of supporting productive antitumor immunity (Eng et al., 2019; Sharma et al., 2023). Experimental studies further suggest that resistance in MSS CRC is not explained simply by a complete absence of tumor antigens; rather, these tumors can harbor clonal neoantigens but fail to support efficient T cell priming, enabling early immune escape (Westcott et al., 2021).

The intestinal microbiota has emerged as an important regulator of antitumor immunity and of response to immune checkpoint blockade immunotherapy in both preclinical and clinical settings (Baruch et al., 2021; Daillere et al., 2016; Davar et al., 2021; Gopalakrishnan et al., 2018; Lee et al., 2022; Matson et al., 2018; McCulloch et al., 2022; Routy et al., 2018; Sivan et al., 2015; Vetizou et al., 2015). Colonization with *Helicobacter hepaticus* was shown to promote CD4+ T follicular helper cell and B cell-dependent antitumor immunity together with the formation of tertiary lymphoid structures, establishing that defined commensal organisms can reshape local immune responses in a model of colitis-associated colon cancer (Overacre-Delgoffe et al., 2021). At the same time, the effects of Helicobacter species are context dependent. In other settings, Helicobacter infection has been linked to chronic intestinal inflammation, mutagenesis, and colorectal tumorigenesis, underscoring that the consequences of these host–microbe interactions depend on species, host immune state, and disease context (Fox et al., 2011; Ge et al., 2019). However, most work in this area has focused on professional antigen-presenting cells, lymphoid organization, or cytokine-dependent effects on the immune compartment. Whether microbiota-derived signals also act directly on malignant epithelial cells in established CRC to alter antigen presentation, metastatic progression, and therapeutic responsiveness remains unclear.

A plausible link between microbial cues and cancer cell immunogenicity is major histocompatibility complex class II (MHC-II). We previously showed that MHC-II expression in intestinal epithelial cells is regulated by a *Helicobacter*-containing gut microbiome and that loss of epithelial MHC-II promotes intestinal tumorigenesis, supporting a role for epithelial antigen presentation in immune surveillance at early stages of disease (Beyaz et al., 2021). In other tumor settings, enforced CIITA-driven MHC-II expression on cancer cells can directly prime naive CD4+ T cells, and cancer cell-intrinsic MHC-II expression has been associated with favorable immune responses and improved outcomes in multiple cancers, including melanoma, lymphoma, colorectal cancer, breast cancer, and more recently, human endometrial cancer (Bou Nasser Eddine et al., 2017; Cho et al., 2021; Chung et al., 2026; Forero et al., 2016; Johnson et al., 2016; Riaz et al., 2017; Rodig et al., 2018; Roemer et al., 2018). At the same time, tumor-cell MHC-II is not uniformly protective. In breast cancer lymph node metastasis, cancer-cell MHC-II promotes regulatory T cell expansion and metastatic progression (Lei et al., 2023). Furthermore, tumor-specific MHC-II can also support adaptive resistance through its function as a ligand for T cell co-inhibitory receptor LAG-3, indicating that the output of this pathway depends on tissue context and immune state (Johnson et al., 2018). Thus, in CRC, it remains unresolved whether microbiota-dependent regulation of epithelial MHC-II persists after malignant transformation, whether cancer-cell MHC-II supports productive immunity or tolerance in established tumors, and whether this axis influences metastatic spread and response to checkpoint blockade immunotherapy.

Here, we identify a microbiome-dependent mechanism that restores MHC-II expression in colon cancer cells and converts immune-refractory MSS CRC into a more immunologically responsive state. Using orthotopic, genetically defined CRC models and autologous human patient-derived organoid – immune co-culture systems, we show that Helicobacter-containing microbiota induce cancer MHC-II expression, which promotes anti-tumor immunity against MSS CRC. Cancer cell-intrinsic MHC-II is required for the antitumor effects of the microbiome and, when enforced, is sufficient to partially phenocopy microbial protection and sensitize MSS tumors to checkpoint blockade immunotherapy. These findings define a microbiome – cancer antigen presentation axis as a mechanistically actionable determinant of immune surveillance and immunotherapy response in MSS CRC.

## RESULTS

### Gut microbiome containing Helicobacter species limits CRC progression and metastasis

To test whether a *Helicobacter*-containing gut microbiome modulates CRC progression, we used an endoscopy-guided orthotopic implantation model that enables longitudinal assessment of tumor growth in the colonic mucosa. Using the MC38 cell line, a hypermutated, MSI-H-like syngeneic CRC model in C57BL/6 mice, we observed a significant reduction in primary tumor burden in *Helicobacter*-colonized (H+) mice compared to *Helicobacter* negative (H-) controls, as measured by colonoscopy and luciferase activity (Figure S1A-E). The incidence of metastasis was also reduced in H+ mice (Figure S1F-G). These anti-tumor effects were reproduced in an independent syngeneic setting using the CT26 tumors in BALB/c mice (Figure S1H-K), indicating that the anti-tumor effects of H+ microbiome generalize across host strains and tumor models.

Because transplanted 2D cell lines do not fully capture the genetics, histopathology and metastatic behavior of human CRC, particularly the predominance of the immune evasive MSS subtype, we next evaluated genetically-defined organoid-based orthotopic CRC models (Roper et al., 2017). Implanting genetically-engineered *Apc^null^, Kras^G12D^, Tp53^null^* (AKP) organoids into the colon leads to formation of tumors that faithfully model MSS CRC (Roper et al., 2017; Westcott et al., 2021). H+ colonization dampened the AKP tumor growth and metastasis (Figure 1A-D, S1L, M), resulting in prolonged overall survival in mice (Figure 1E).

**Figure 1.**
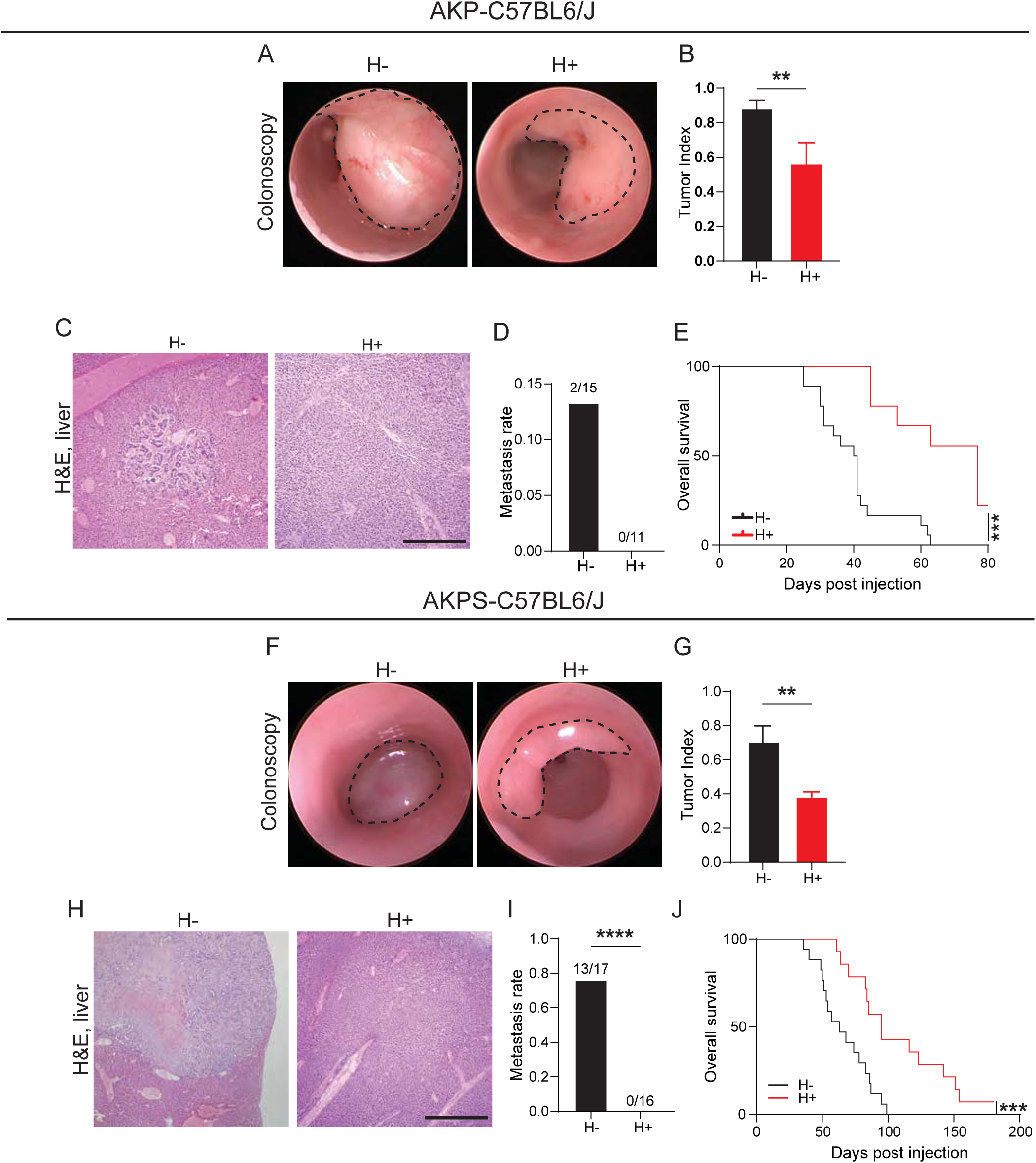
Helicobacter containing gut microbiome regulates primary tumor progression and metastasis in orthotopic MSS CRC models. (A) Representative colonoscopy images of colon tumors induced by AKP organoids in H- and H+ mice at 5 weeks after AKP organoid implantation. (B) Tumor index measured 5 weeks after AKP organoid implantation (H-, n=9; H+, n=7). (C) Representative H&E staining of liver metastases arising from AKP-induced primary tumors in H- and H+ mice. Scale bar, 500μm. (D) Frequency of liver metastasis in mice bearing AKP-induced tumors (H-, n=15; H+, n=11). (E) Overall survival of H- and H+ mice with AKP-induced tumor (H-; n=15, H+; n=9). Statistical comparisons were performed using Kaplan-Meier analysis (*** p<0.001). (F) Representative colonoscopy images of colon tumors induced by AKPS organoids in H- and H+ mice at 6 weeks after AKPS organoid implantation. (G) Tumor index measured 6 weeks after AKPS organoid implantation (H-n=7; H+, n=9). (H) Representative H&E staining of liver metastasis arising from AKPS-induced tumors in H-and H+ mice. (I) Frequency of liver metastasis in mice bearing AKPS-induced tumors (H-, n=17; H+, n=16). Scale bar, 500μm. (J) Overall survival rate of H- and H+ mice with AKPS-induced tumors (H-, n=17; H+, n=16). Statistical comparisons were performed using Kaplan-Meier analysis was (*** <0.001). Unpaired T-test were used for tumor index comparisons in (B) and (G). Fisher’s exact test was used for metastasis frequency in (D) and (I) (**** p<0.0001).

We next asked whether *Helicobacter* colonization could constrain a more aggressive and immune evasive MSS model with enhanced metastatic competence. Mice implanted orthotopically with genetically engineered AKPS (*Apc^Knock-down^*, *Kras^G12D^*, *Tp53^null^*, *Smad4^null^*) organoids, develop aggressive and immune-evasive MSS tumors that eventually metastasize to the liver and other sites at enhanced frequency (Roper et al., 2017; Westcott et al., 2021). In this model, we found that *Helicobacter* colonization restricts primary tumor growth (Figure 1F-G), and strikingly, prevented metastasis (Figure 1H-I), leading to significant survival advantage in H+ mice (Figure 1J). Together, these data identify a Helicobacter-associated gut microbiome flora that restricts both tumor progression and metastasis across four different CRC models spanning distinct genetic and immunologic contexts in two different mouse strains.

### Colonization with single Helicobacter species partially restricts CRC progression and metastasis

We previously found that H+ community contain Helicobacter species *H. typhlonius* or *H. mastomyrinus* (Beyaz et al., 2021). To determine whether individual *Helicobacter* species identified in the H+ community are sufficient to restrain tumor progression, we inoculated H-mice with either *H. typhlonius* or *H. mastomyrinus*, confirmed successful colonization (Figure S2A–G), and then orthotopically implanted AKPS organoids. Colonization with either species reduced primary tumor burden and metastatic incidence (Figure S2H–J) and improved survival relative to H-controls (Figure S2K). However, these effects were consistently less pronounced than those observed following whole-flora transfer. These findings indicate that single *Helicobacter* species can confer partial protection while additional microbial constituents or cooperative community interactions may be needed for the full protective effects of whole-flora transfer.

To determine whether the anti-metastatic activity we observed reflects a broader genus-level property of *Helicobacter*, we evaluated additional *Helicobacter* species that were not detected in the H+ flora. We inoculated H-mice with *H. bilis*, *H. pullorum*, or *H. hepaticus* and assessed metastatic dissemination following orthotopic implantation of AKPS tumors. In contrast to H+ flora, colonization with these species did not reduce metastatic incidence (Figure S2L). Together, these data indicate that suppression of metastatic progression is not a universal feature of *Helicobacter* colonization in mice.

### Helicobacter colonization enhances intratumoral lymphocyte infiltration and effector programs

Tumor-infiltrating lymphocytes are strongly associated with favorable outcomes in CRC (Galon et al., 2006), and defined commensals can augment intratumoral immune infiltration to promote anti-tumor immunity (Sivan et al., 2015; Vetizou et al., 2015). Consistent with the improved survival observed in H+ animals, Helicobacter colonization increased intratumoral CD8+ T cell infiltration in the AKPS model, as assessed by immunofluorescence (Figure 2A, B). We observed a similar increase in CD8+ T cell accumulation in the more immunogenic MC38 model (Figure S3A, B), indicating that Helicobacter-associated microbiota promotes T cell infiltration across CRC models of both MSS and MSI-H disease.

**Figure 2.**
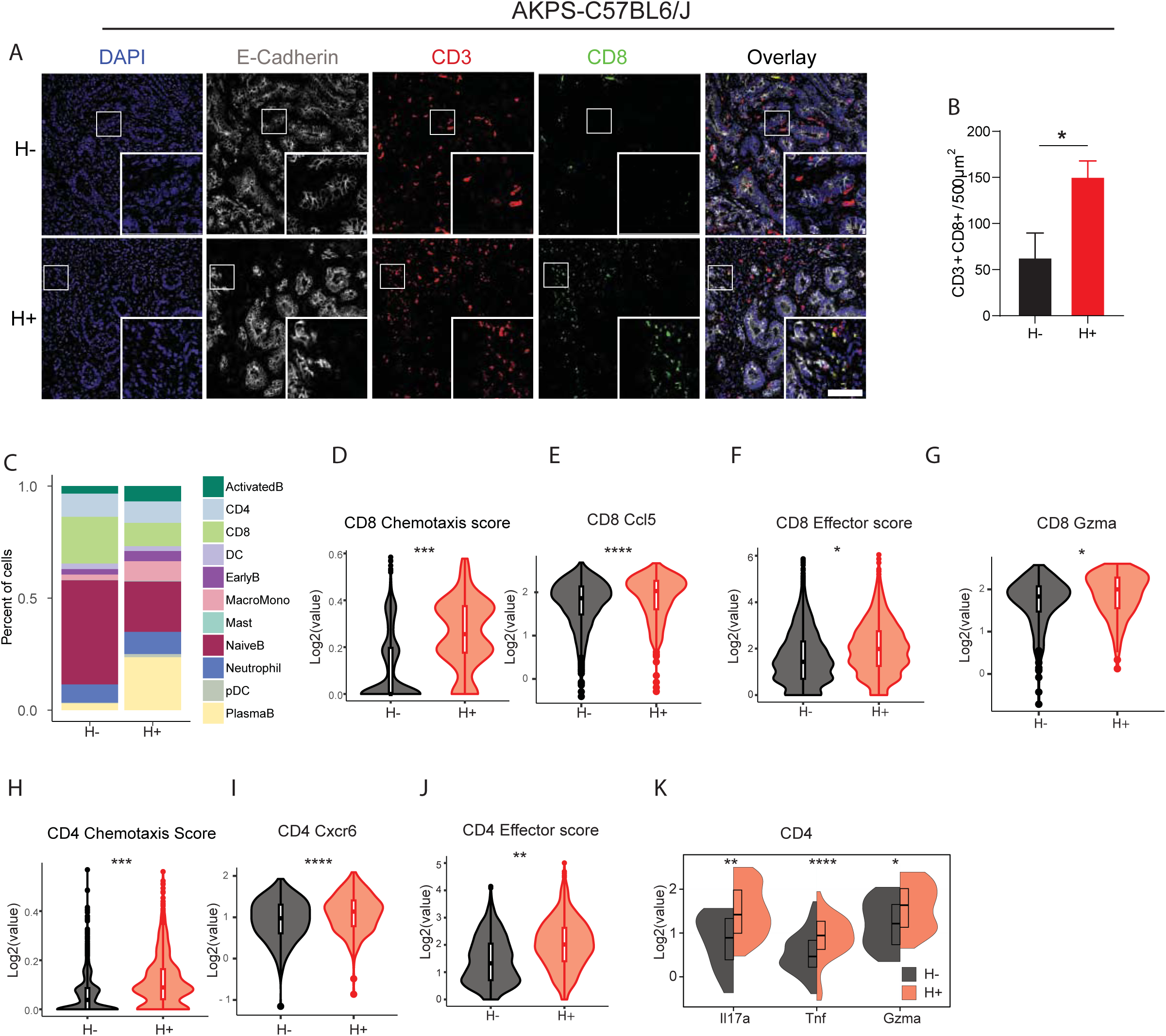
Helicobacter containing gut microbiome promotes TIL recruitment and enhance T-cell effector programs in MSS CRC. (A) Representative immunofluorescence (IF) images of AKPS Tumors stained for DAPI (blue), E-cadherin (white), CD3 (red), and CD8 (green), illustrating spatial organization of T cells relative to epithelial structures. Scale bars: 50μm (low magnification), 12.5μm (high magnification). (B) Quantification of CD8^+^ T cells (CD3^+^ CD8^+^) co-localized with EpCAM^+^ tumor glands in tumors from H- and H+ mice (H-, n=4; H+, n=5). (C) Proportions of clustered CD45^+^ cells from digested AKPS tumors from H- or H+ mice (n=4 per group). (D-G) Violin plots showing chemotaxis and effector programs in CD8^+^ cell. Chemotaxis score (D), Ccl5 expression (E), effector score (F), and *Gzma* expression (G). (H-K) Violin plots showing chemotaxis and effector programs in CD4^+^ T cells. Chemotaxis score (H), *Cxcr6* expression (I), effector score (J), and expression of effector cytokines *Il17a*, *Tnf*, and *Gzma* (K). Unpaired *t*-tests were used for statistical comparisons (B, D-K). Gene sets used to compute chemotaxis and effector scores are listed in supplementary table 2.

To define how Helicobacter colonization remodels the tumor-immune microenvironment at higher resolution, we performed single-cell RNA sequencing on FACS-sorted CD45+ and CD45-fractions from enzymatically dissociated AKPS tumors harvested 21 days after orthotopic implantation in H+ and H-mice. After quality control, we analyzed the CD45+ compartment by unsupervised clustering and annotated immune populations using canonical lineage markers (Figure S4A, B). The overall representation of major immune lineages was broadly similar between groups. However, H+ tumors showed shifts in composition, including increased macrophage and monocyte populations and enrichment of B lineage states, most prominently plasma B cells (Figure 2C). Within the T cell compartment, Helicobacter colonization was associated with coordinated induction of trafficking and effector programs. Intratumoral CD8+ T cells from H+ tumors showed higher chemotaxis signatures and increased expression of *Ccl5* (Figure 2D, E), a chemokine implicated in tumor T cell infiltration programs (Topper et al., 2023). CD8+ T cells in H+ tumors also exhibited increased effector scores together with higher expression of *Gzma* (Figure 2F, G), consistent with enhanced cytotoxic differentiation. CD4+ T cells similarly demonstrated increased chemotaxis signatures and higher *Cxcr6* expression (Figure 2H, I), a chemokine receptor linked to intratumoral localization and persistence of T cells (Di Pilato et al., 2021). This was accompanied by increased CD4 effector scores and higher expression of inflammatory and cytotoxic transcripts including *Il17a*, *Tnf*, and *Gzma* (Figure 2J, K).

We next used the MC38 model as an orthogonal system to test whether the immune-state shifts observed in AKPS tumors were conserved. MC38 scRNA-seq recapitulated key features observed in AKPS, with H+ tumors enriched for plasmablasts (Figure S4C–E) and T cells skewed toward effector differentiation (Figure S5A-C). Specifically, cytotoxic CD8+ T cells exhibited increased cytotoxicity gene scores with higher *Gzma*/*Gzmb* and reduced exhaustion-associated *Pdcd1*/*Lag3* (Figure S5A–C) expression. CD4+ T cells showed increased levels of effector-associated transcripts (*Il18r1*, *Jun*, *Slamf6*) along with higher *Cxcr6* expression (Figure S5D). These findings demonstrate that Helicobacter colonization is associated with conserved enhancement of intratumoral T cell infiltration and effector programs across CRC models.

### T cells, B cells and Cxcr6+ cells are required for microbiome-mediated tumor control

Single-cell RNA-seq analyses revealed enhanced chemotactic and effector programs in CD4^+^ and CD8^+^ T cells from H+ tumors, accompanied by an expansion of mature B-cell populations, including plasma B cells, in both AKPS and MC38 models. These observations raised the possibility that adaptive immune cells contribute directly to microbiome-associated restriction of tumor progression and metastasis. To determine whether adaptive lymphocytes are required for microbiome-mediated tumor control, we depleted T cells (CD4^+^ and CD8^+^) or B cells (CD19^+^ and B220^+^) in H+ mice bearing AKPS tumors. T cell depletion resulted in a marked increase in metastatic incidence and significantly reduced survival compared with non-depleted H+ controls (Figure 3A, B). Similarly, depletion of B cells increased metastatic burden and shortened survival relative to H+ controls (Figure 3A, B). Together, these results indicate that both T cells and B cells contribute to microbiome-associated restriction of metastatic progression.

**Figure 3.**
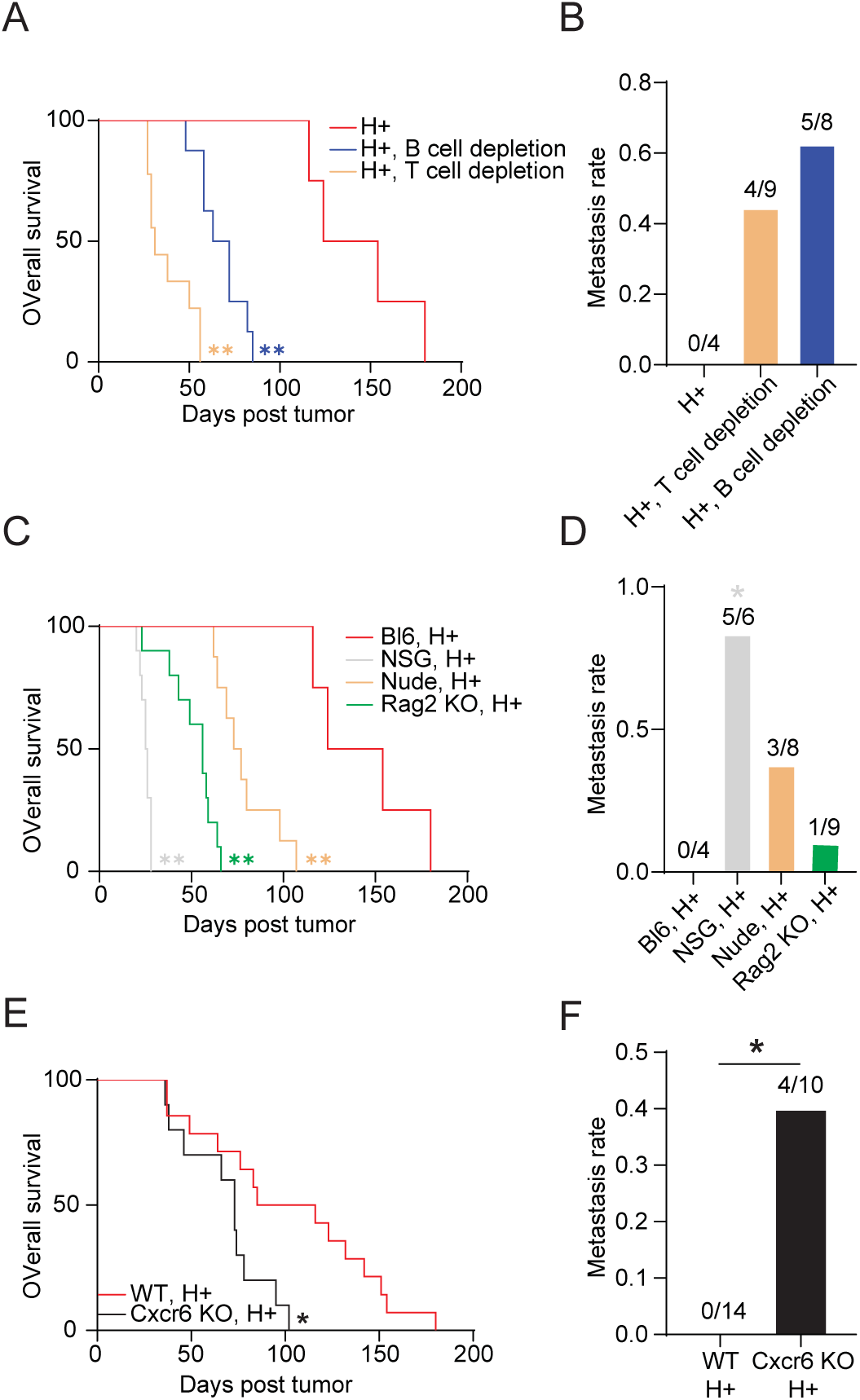
Microbial anti-tumor effects require T, B and Cxcr6+ cells. (A) Overall survival of helicobacter-colonized C57BL/6 mice bearing AKPS tumors following B-cell or T cell depletion (H+, n=4; B cell depletion, n=8; T cell depletion, n=9). B cells (anti-CD19 + anti-B220) and T cells (anti-CD4 + anti-CD8) were depleted by antibody administration as described in Methods. Kaplan-Meier analysis was used for statistical comparisons (** p<0.01). (B) Liver metastasis rate in helicobacter-colonized, B-cell or T-cell depleted, C57BL/6 mice with AKPS tumors (H+, n=4; B cell depletion, n=8; T cell depletion, n=9). (C) Overall survival of helicobacter-colonized NSG, Nude, Rag2 KO, and C57BL/6 mice with AKPS tumors (H+, n=4; NSG, n=6; Nude, n=8; Rag2 KO, n=9). Kaplan-Meier analysis was used for statistical comparisons (** p<0.01). (D) Liver metastasis rate in helicobacter-colonized NSG, Nude, Rag2 KO, and C57BL/6 mice with AKPS-induced tumor (H+, n=4; NSG, n=6; Nude, n=8; rRag2 KO, n=9). (E) Overall survival of helicobacter-colonized C57BL/6and Cxcr6 KO mice with AKPS-induced tumors (H+, n=14; Cxcr6 KO, n=10). Kaplan-Meier analysis (* p<0.05). (F) Liver metastasis rate in helicobacter-colonized C57BL/6and Cxcr6 KO mice with AKPS-induced tumor (H+, n=14; Cxcr6 KO, n=10). Fisher’s exact test was used for statistical comparisons (B, D, and F).

To further assess the role of adaptive immunity, we evaluated tumor progression and metastasis in immunodeficient mouse models under H+ conditions, including NSG (lacking T, B and NK cells), Nude (lacking T cells), and Rag2 KO (lacking both T and B cells) mice. Loss of adaptive immune components diminished the metastasis control and survival benefit of the H+ flora across these models (Figure 3C-D). The persistent anti-metastatic effect in Rag2 KO mice likely reflects a preserved NK-cell mediated surveillance against metastasis (Figure 3D).

In parallel, our scRNA-seq analysis identified consistently elevated *Cxcr6* expression in CD4^+^ T cells from H+ tumors across both AKPS and MC38 models (Figure 2I, Figure S5D). Given the established association of Cxcr6 with tissue-resident memory T-cell positioning within tumors (Di Pilato et al., 2021), we tested its functional relevance by implanting the AKPS organoids into Cxcr6-deficient mice under H+ conditions. Cxcr6 deficiency resulted in increased metastatic incidence and reduced survival compared to wild-type controls (Figure 3E-F), indicating that Cxcr6+ cells contribute to microbiome-mediated restriction of tumor progression and metastasis. Collectively, these results demonstrate that the protective effects of H+ microbiota require coordinated adaptive immune responses, including both T cells and B cells, and tissue-resident memory-associated Cxcr6+ cells.

### Helicobacter colonization enhances T cell priming

Because effective tumor infiltration and cytotoxic activity require proper priming of T cells within tumor-draining lymph nodes (dLN), we next asked whether Helicobacter colonization enhances tumor-specific T-cell priming. To directly assess priming potential of CD8^+^ T cells, we implanted AKPS organoids expressing low levels of the ovalbumin peptide SIINFEKL (AKPS loSIIN (Westcott et al., 2021)) into H- and H+ mice, with microbial colonization established 7 days prior to tumor implantation. T cell responses were analyzed by flow cytometry at days 7 and 14 following tumor implantation. At an early time point (day 7), H+ mice exhibited a pronounced expansion of CD8^+^ T cells in the dLN, accompanied by increased numbers of CD4^+^ T cells, consistent with enhanced priming in secondary lymphoid organs (Figure S6A-B). This early expansion preceded a significant increase in CD8^+^ T cell accumulation within tumors at day 14 (Figure S6C-D), indicating a temporal progression from lymph node priming to intratumoral effector accumulation.

Because the magnitude of effector T cell priming is reflected by the expansion of antigen-specific T cells expressing cytokines and cytolytic molecules, including granzyme B (GzmB), IFNγ *and* TNFα, (Westcott et al., 2021), we next assessed functional differentiation of T cells in AKPS tumor bearing mice. In the dLN, H+ tumor-bearing mice displayed increased numbers of antigen-specific CD8^+^ T cells expressing Gzmb, IFNγ (Figure 4A-B), and TNFα (Figure S6E) at both time points, days 7 and 14. Similarly, CD4^+^ T cells expressing GzmB, IFNγ (Figure 4C-D), and TNFα (Figure S6F) were enriched in the dLN of H+ tumor bearing mice. In agreement with this early priming phenotype, H+ mice exhibited increased accumulation of cytotoxic tumor-specific CD8^+^ T cells within tumors. Consistent with the effector programs observed in the draining lymph nodes, antigen-specific CD8^+^ T cells within tumors of H+ mice also exhibited increased expression of Gzmb, IFNγ (Figure 4E-F) and TNFα (Figure S6G) at both days 7 and 14. Notably, CD4^+^ T cells expressing GzmB, IFNγ (Figure 4G-H) and TNFα (Figure S6H) were also significantly enriched in H+ tumors, indicating induction of a cytotoxic transcriptional program within the CD4^+^ T-cell compartment in the context of Helicobacter colonization.

**Figure 4.**
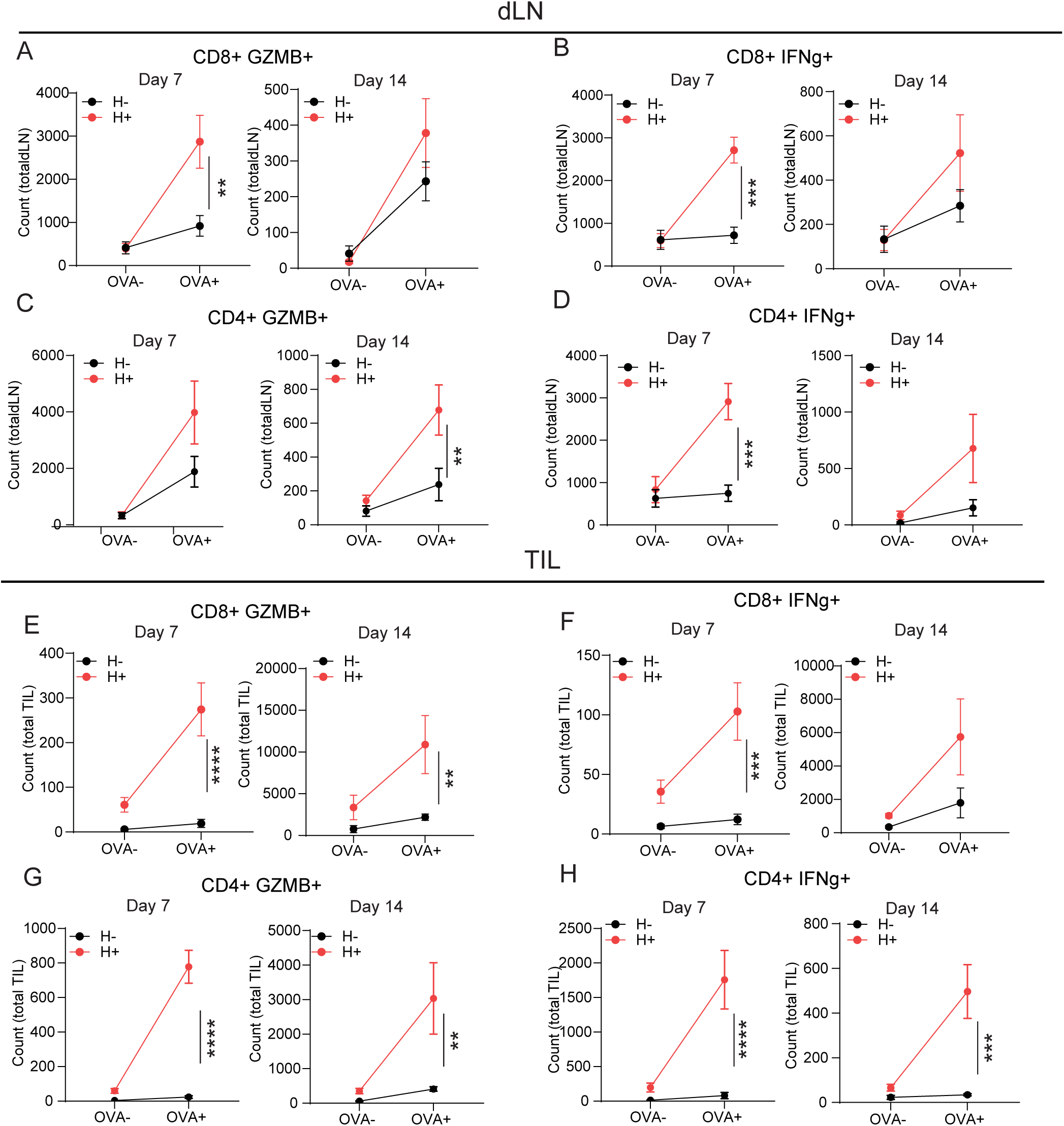
Helicobacter containing gut microbiome enhances T cell priming. (A-D) Quantification of GzmB^+^ or IFNγ^+^ tumor-specific (SINFEKL+) CD8+ and CD4+ T cells in the draining lymph nodes of AKPS tumor-bearing H- and H+ mice on day 7 (n=5 per group) and day 14 (H-, n=8; H+; n=9), with or without OVA (SINFEKL) stimulation. (E-H) Quantification of GzmB^+^ or IFNγ^+^ tumor specific (SINFKL+) CD8+ and CD4+ T cells infiltrating AKPS tumors from H- and H+ mice on day 7 (n=5 per group) and day 14 (H-, n=8; H+, n=9), with or without antigen restimulation. Unpaired *t*-test was used to assess statistical significance (* p<0.05, ** p<0.01, *** p<0.001, **** p<0.0001).

### Tumor MHC-II expression is necessary and sufficient for the microbiome-mediated anti-tumor effects

We next asked whether the presence of *Helicobacter* species in the microbiome modulates MHC-II expression on cancer cells. To address this, we revisited our scRNA-seq dataset and examined non-immune (CD45^-^) clusters within both AKPS and MC38 tumors (Figure S7A-D). Tumor cells from H+ mice in both models exhibited significant increase in MHC associated genes (Figure 5A-B; Figure S7E-H), which was mirrored by elevated MHC protein levels (Figure 5 C-D; Figure S7I-J).

**Figure 5.**
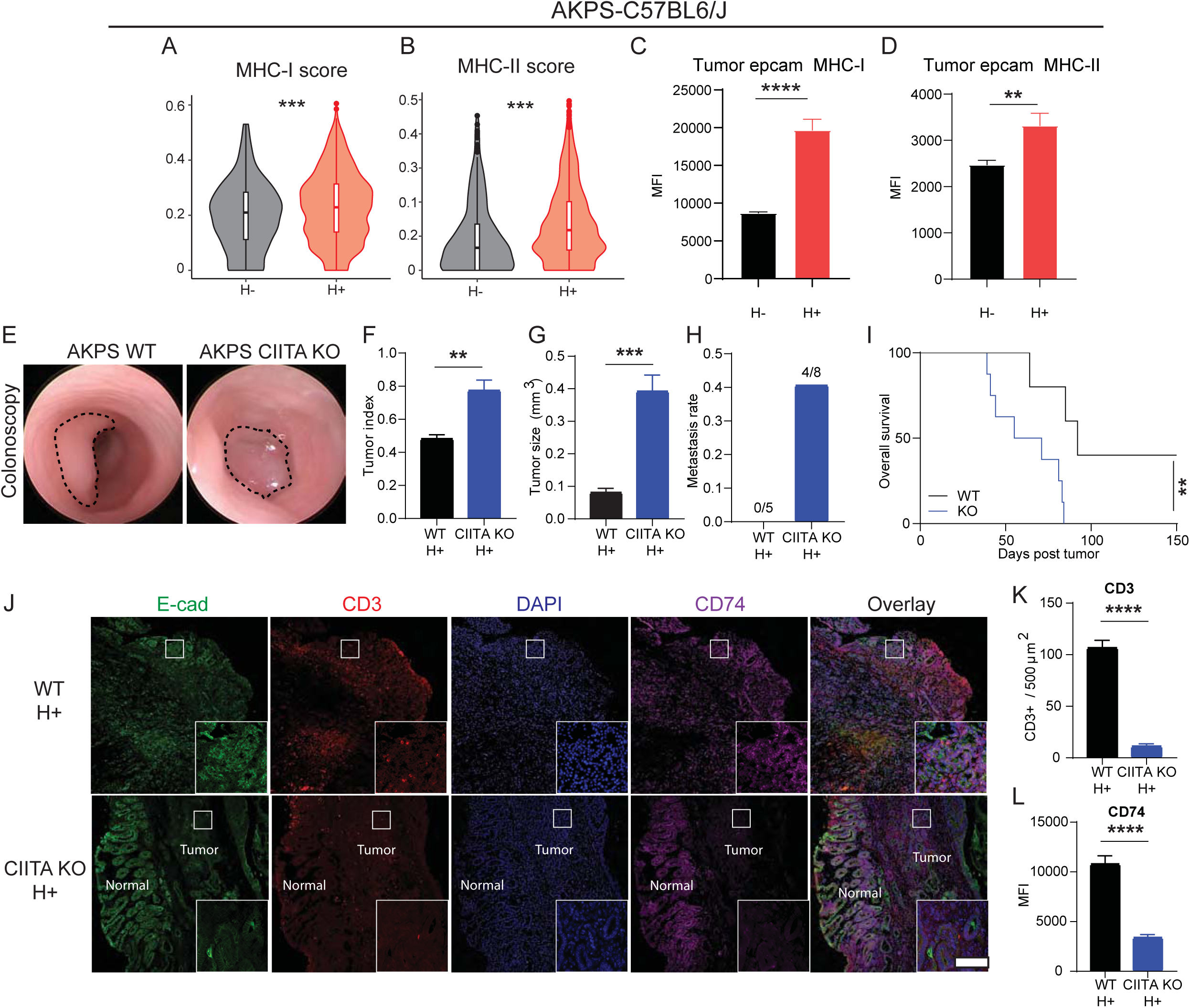
Cancer cell-intrinsic MHC-II expression is required for microbe-driven anti-tumor immunity against MSS CRC. (A-B) Violin plots of MHC-I (A) and MHC-II (B) gene score in cancer cell subclusters from CD45^-^cells isolated from digested AKPS tumors (n=4 per group) (C-D) MHC-I (C) and MHC-II (D) expression levels of EpCAM^+^ cells from digested AKPS tumors measured by flow cytometry (n=4 per group) (E) Representative colonoscopy images of colon tumors induced by AKPS WT or AKPS CIITA KO organoids in H+ mice at day 15. (F) Tumor index of AKPS WT or AKPS CIITA KO organoid induced tumors in H+ mice at day 15 (WT, n=5; CIITA OE, n=8). (G) Tumor size measured at day 21 following implantation (n=4 per group). (H) Liver metastasis rates in H+ mice bearing AKPS WT or AKPS CIITA KO tumors (WT, n=5; CIIAT KO, n=8). (I) Overall survival rate of H+ mice bearing either AKPS WT or AKPS CIITA KO induced tumors (WT, n=5; CIITA KO, n=8). Kaplan-Meier analysis was used for survival comparisons (** p<0.01). (J) Representative immunofluorescence (IF) images of tumors induced by AKPS WT or AKPS CIITA KO organoids. Scale bar, 500μm. (K) Quantification of CD74 by mean immunofluorescence intensity (MFI) in E-cadherin^+^ tumor epithelia regions (WT, n=4; KO, n=4) (L) Quantification of CD3^+^ T cells within tumor regions (WT, n=4; KO, n=4). Unpaired *t*-tests were used for statistical analysis (A-D, E-F, K-L) and Fisher’s exact test was used for metastasis frequency analysis (H). (** p<0.01, *** p<0.001, **** p<0.0001).

Next, we tested whether tumor-intrinsic MHC-II mediates microbial anti-tumor immunity. To selectively ablate tumor MHC-II while preserving MHC-I upregulation, we generated a CIITA deficient AKPS organoid (CIITA KO). As expected, CIITA KO organoids upregulated MHC-I, but not MHC-II, in response to IFNγ (Figure S8A-B). Implantation of CIITA KO tumors into H+ mice resulted in increased tumor volume (Figure 5E-G), increased metastasis (Figure 5H), and reduced survival (Figure 5I) compared to parental tumors. Immunofluorescence analysis demonstrated reduced CD3^+^ T cell infiltration in CIITA KO tumors (Figure 5J-K), accompanied by loss of tumor-cell MHC-II pathway activity as reflected by reduced CD74 staining (Figure 5L). Together these finding indicate that tumor-intrinsic MHC-II is required for full microbiome-mediated anti-tumor immunity.

Building on the requirement for tumor cell-intrinsic MHC-II in the H+ setting, we next asked whether tumor MHC-II alone is sufficient to restrict tumor progression and metastasis, in the absence of microbial colonization. To this end, we generated an AKPS organoid with elevated CIITA expression (CIITA OE). The CIITA OE AKPS organoids displayed significantly high MHC-II expression in the absence of external stimuli, such as IFNγ (Figure S8C-D). Using CIITA OE and parental AKPS organoids, we assessed whether enforced tumor-intrinsic MHC-II expression confers anti-tumor activity in H-mice. Our results demonstrated that MHC-II overexpression significantly reduced tumor volume (Figure 6A-C) and metastasis rate (Figure 6D). However these reductions did not result in significant improvement in overall survival (Figure 6E), indicating that tumor MHC-II is sufficient to partially restrict tumor progression and metastasis but does not fully recapitulate the survival benefit associated with H+ microbiome colonization. Tumors derived from CIITA OE organoids contained increased numbers of intratumoral T cells by immunofluorescence analysis (Figure 6F-G) alongside heightened tumor-cell MHC-II expression as reflected by CD74 staining (Figure 6H). Collectively, these data demonstrate that tumor-intrinsic MHC-II expression promotes intratumoral T-cell presence and contributes to restriction of tumor growth and metastasis, and that enforced MHC-II expression on cancer cells is sufficient to confer partial anti-tumor activity even in the absence of Helicobacter colonization.

**Figure 6.**
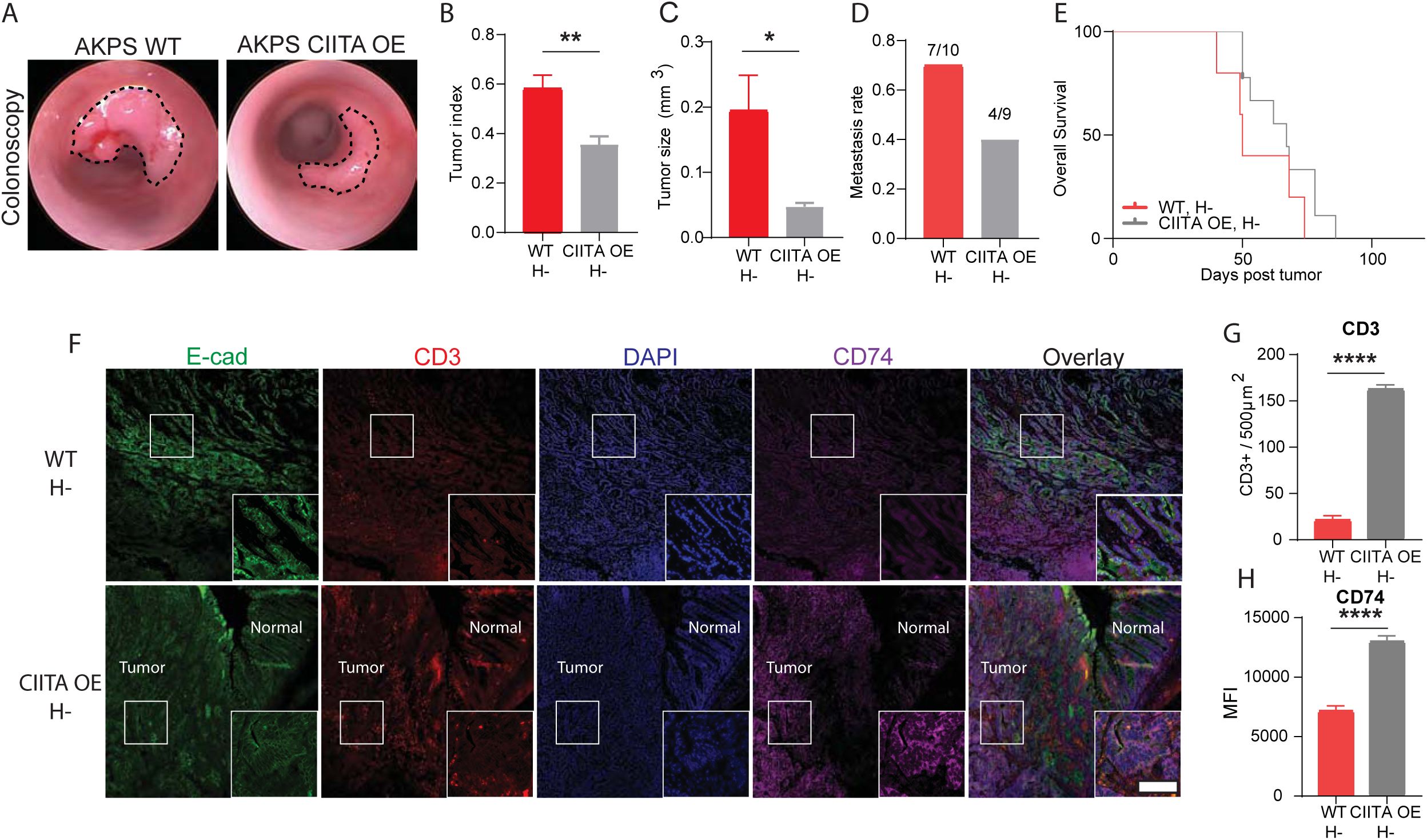
Cancer cell-intrinsic MHC-II expression is sufficient to elicit anti-tumor immunity against MSS CRC. (A) Representative colonoscopy images of colon tumors induced by AKPS WT or AKPS CIITA OE organoids in H-mice at day 15. (B) Tumor index of AKPS WT or AKPS CIITA OE organoid induced tumors in H-mice at day 15 (WT, n=5; CIITA OE, n=9). (C) Tumor size measured at day 21 following implantation (n=4 per group). (D) Liver metastasis rate of H-mice bearing AKPS WT or AKPS CIITA OE tumors (WT, n=9; CIIAT OE, n=9). (E) Overall survival of H-mice bearing AKPS WT or AKPS CIITA OE tumors (n=9 per group). (F) Representative immunofluorescence (IF) images of tumors induced by AKPS WT or AKPS CIITA OE organoids. Scale bar, 500μm. (G) Quantification of CD74 mean immunofluorescence intensity (MFI) in EpCAM^+^ tumor epithelia regions (WT, n=4; CIITA OE, n=4). (H) Quantification of CD3^+^ T cells within the tumor regions (WT, n=4; CIITA OE, n=4). Unpaired *t*-tests were used for statistical analyses (B-C and G-H) and Fisher’s exact test was used for metastasis frequency (D). (* p<0.05, ** p<0.01, **** p<0.0001).

### MHC-II mediated anti-tumor effects require immune cell trafficking from the lymph node

Our data show that microbiome-induced upregulation of tumor MHC-II enhances T-cell priming, increases tumor-infiltrating lymphocytes, reduces tumor burden and decreases metastasis incidence. These findings may suggest that anti-tumor effects of cancer-cell MHC-II depend on effective immune cell priming within tumor-draining lymph nodes and subsequent trafficking into the tumor microenvironment. To test this, we implanted either WT or CIITA-overexpressing (CIITA OE) AKPS organoids and treated mice with vehicle or FTY720, an inhibitor of lymphocyte egress from lymph nodes. Tumor burden in CIITA OE tumors treated with FTY720 was comparable to CIITA WT controls, indicating that elevated tumor-cell MHC-II alone is not sufficient to restrict tumor growth when the trafficking of immune cells from the lymph node is inhibited (Figure S8E-H).

### Helicobacter colonization after tumor implantation restricts metastasis

Inspired by the microbiome-mediated anti-tumor effects observed in our preclinical models, we next assessed whether *Helicobacter* colonization could exert therapeutic benefit when introduced after tumor establishment. AKPS organoids were implanted into H-mice, and once tumors were confirmed at day 7, half of the mice were subsequently inoculated with H+ flora. While H+ colonization after tumor implantation slowed tumor growth by 3 weeks compared to controls, it did not reduce tumor burden at endpoint (Figure S9A-D) but resulted in a significant reduction in metastasis incidence (Figure S9E) and improved metastasis-free survival (Figure S9F) although overall survival was not significantly altered (Figure S9G). These data indicate that *Helicobacter* colonization after tumor establishment can restrict metastasis.

### Augmenting cancer MHC-II expression enhances immunotherapy efficacy in MSS CRC

We next asked whether modulation of tumor-intrinsic MHC-II expression, induced either through *Helicobacter* colonization or CIITA overexpression, could enhance the response of MSS tumors to immune-checkpoint blockade (ICB, anti-PD-1 + anti-CTLA-4; Figure 7A). As expected, ICB alone did not control tumor growth in H-mice. In contrast, when ICB was combined with post-tumor *Helicobacter* inoculation, tumor growth was significantly reduced by the end of treatment (Figure 7B-D). Similarly, the growth of CIITA-overexpressing tumors were restricted in response to ICB compared to controls (Figure 7B-C), indicating that tumor-intrinsic MHC-II increases sensitivity of MSS tumors to ICB. Complete tumor rejection was rare in control animals (10%, 1/10) in the H-group, whereas the rejection rate was increased to 30% (3/10) in H+ mice and 71% (5/7) in mice bearing CIITA-overexpressing tumors (Figure 7D). Consistent with this, *Helicobacter* colonization or CIITA overexpression significantly reduced the incidence of metastasis compared to the controls (Figure 7E) and improved survival (Figure 7F). Collectively, these findings demonstrate that elevating cancer MHC-II expression augments the response to immunotherapy.

**Figure 7.**
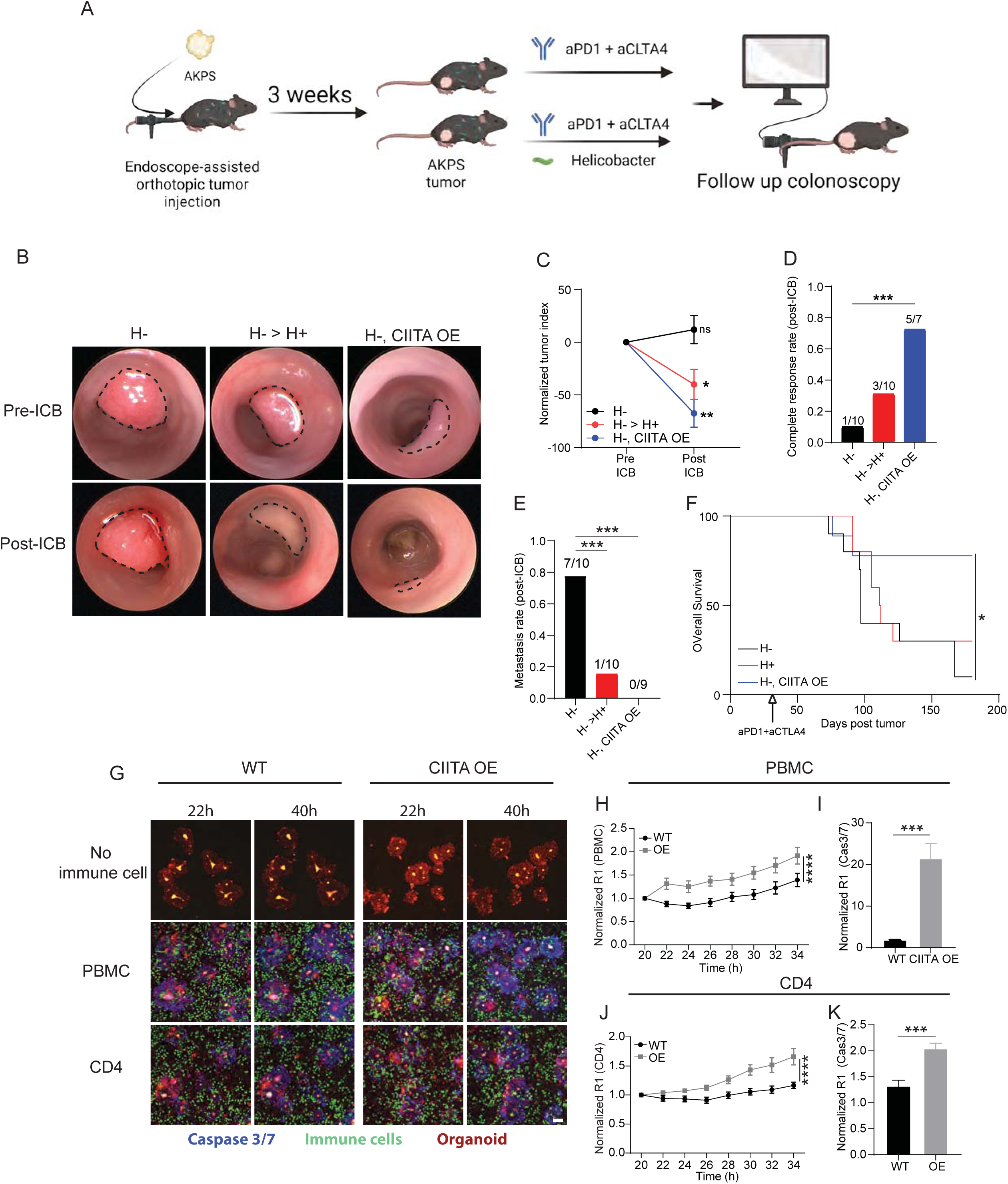
Cancer MHC-II synergizes with ICB therapy and augments immune response in human autologous CRC PDO - immune cell co-culture system. (A) A schematic depicting post-tumor implantation immunotherapy experimental design. (B) Representative colonoscopy images of colon tumors induced by AKPS WT in H-, post-tumor H+, and AKPS CIITA OE in H-mice on day 21 (pre-ICB) and 35 (post-ICB). Post-tumor inoculation and immunotherapy was concomitantly initiated on day 7. (C) Normalized tumor index of colon tumors induced by AKPS WT in H-, post-tumor H+, and AKPS CIITA OE in H-mice on day 21 (pre-ICB) and 35 (post-ICB). (H-; n=10, H--> H+; n=10, CIITA OE; n=7). Tumor index was normalized to pre-treatment measurement for respective groups. (D) Complete response rate upon immunotherapy of the mice in (C) by day 35. (H-; n=10, H--> H+; n=10, H-CIITA OE; n=7) (E) Rate of metastasis of H- and H--> H+ mice bearing AKPS WT or AKPS CIITA OE induced tumors with immunotherapy (H-; n=10, H--> H+; n=10, H-CIITA OE; n=7). (F) Overall survival of H- and H--> H+ mice bearing AKPS WT or AKPS CIITA OE induced tumors with immunotherapy (H-; n=10, H--> H+; n=10, H-CIITA OE; n=9). Kaplan-Meier analysis was applied to calculate statistical significance (* p<0.05) (G) Representative images of co-culture using a MSS CRC PDO line with autologous PBMC or CD4 T cells. Red and green fluorescence represent PDOs and corresponding immune cells, respectively. Blue fluorescence represents active caspase 3/7 signals. Scale bar = 50 µm. (H) Co-localization of the PDOs and PBMCs over the time of imaging. The area under the curve was used for statistical significance (unpaired T-test). (I) End point viability of the PDOs and PBMCs measured by co-localization coefficient between PDOs and caspase 3/7 signals. (J) Co-localization of the PDOs and CD4+ T cells over the time of imaging. The area under the curve was used for statistical significance (unpaired T-test). (K) End point viability of the PDOs and CD4+ T cells measured by co-localization coefficient between PDOs and caspase 3/7 signals. Unpaired *t*-test (C, I, K), and fisher’s exact test (D-E) were used to calculate statistical significance (* p<0.05, ** p<0.01, *** p<0.01)

### Augmenting cancer MHC-II expression enhances anti-tumor immunity in autologous human CRC PDO – immune co-culture system

Finally, we extended these findings to human MSS CRC using a 3 dimensional (3D) autologous patient-derived organoid (PDO) – immune cell co-culture system. CIITA overexpression in a MSS CRC PDO line robustly increased MHC-II expression levels compared to parental WT PDO line (Figure S10A). Over a 3-day co-culture period, autologous PBMCs exhibited minimal cytotoxicity toward WT PDOs but efficiently killed CIITA-overexpressing PDOs with elevated MHC-II expression. To visualize these interactions in real time, we established a confocal live-cell imaging assay using human PDOs co-cultured with autologous PBMCs or autologous CD4+ T cells (Figure 7G, Figure S10B). Increased tumor-cell MHC-II expression in CIITA-overexpressing PDOs enhanced PBMC colocalization with cancer cells (Figure 7H) and increased apoptosis in PDOs engaged by immune cells (Figure 7I). CIITA overexpression similarly increased interactions between PDOs and autologous CD4+ T cells and promoted apoptosis in cancer cells, although these effects were more modest than those observed with PBMCs (Figure 7J-K). Together, these findings indicate that increasing cancer-cell MHC-II expression enhances antitumor immune engagement in human MSS CRC.

## DISCUSSION

Most deaths from CRC result from metastatic progression, and this clinical burden falls predominantly on patients with MSS disease, a setting in which checkpoint blockade immunotherapy has shown limited benefit (Canellas-Socias et al., 2024; Dekker et al., 2019; Eng et al., 2019; Le et al., 2015). In parallel, studies in human CRC and mouse models suggest that MSS tumors are not uniformly invisible to the immune system, but instead many fail at an early step of productive T cell priming and sustained local immune engagement (Galon et al., 2006; Mlecnik et al., 2016; Westcott et al., 2021). Our study places a microbiome-regulated epithelial antigen-presentation program into this framework by showing that a Helicobacter-containing microbiome induces MHC-II expression in colon cancer cells, enhances intratumoral T cell accumulation, restrains tumor progression and metastasis, and sensitizes MSS CRC to immune checkpoint blockade immunotherapy. These findings extend the current models of microbiota-dependent antitumor immunity beyond effects on T cells, professional antigen-presenting cells, cytokine milieu, systemic anti-tumor immunity and tertiary lymphoid structures (Baruch et al., 2021; Daillere et al., 2016; Davar et al., 2021; McCulloch et al., 2022; Overacre-Delgoffe et al., 2021; Routy et al., 2018; Sivan et al., 2015; Vetizou et al., 2015). Our results add a complementary mechanistic layer by showing that microbial signals can act directly on the malignant epithelial compartment, converting tumor cells into putative antigen presenting cells with MHC-II expression.

A central implication of our work is that cancer MHC-II is not simply a passive correlate of inflammatory cytokine exposure, but a causal determinant of antitumor immunity in MSS CRC (Axelrod et al., 2019; Bou Nasser Eddine et al., 2017; Cho et al., 2021; Johnson et al., 2016; Rodig et al., 2018). In multiple tumor types, including melanoma, lymphoma, breast cancer, and endometrial cancer, tumor-intrinsic MHC-II expression has been associated with favorable immune states, improved clinical outcomes, or increased responsiveness to immunotherapy (Bou Nasser Eddine et al., 2017; Cho et al., 2021; Chung et al., 2026; Forero et al., 2016; Rodig et al., 2018). Experimental studies further support functional significance, showing that MHC-II expression on tumor cells promote anti-tumor immunity, prime naive CD4+ T cells, and that MHC-II-restricted neoantigens can shape tumor immunity and response to immunotherapy (Alspach et al., 2019; Beyaz et al., 2021; Bou Nasser Eddine et al., 2017; Kalaora et al., 2021; Marty Pyke et al., 2018). Consistent with these observations, we find here that deletion of CIITA in tumor cells blunts microbe-driven enhancement of anti-tumor immunity, whereas enforced CIITA expression increases intratumoral T cell accumulation, reduces metastasis, and improves sensitivity to checkpoint blockade immunotherapy even in the absence of the protective microbiome state.

At the same time, our data reinforce the idea that the functional consequences of tumor-cell MHC-II expression are context dependent rather than uniformly beneficial across tissues and disease states (Axelrod et al., 2019). In breast cancer metastasis to lymph nodes, MHC-II-expressing cancer cells can promote immune tolerance and regulatory T cell expansion, indicating that the consequences of tumor-intrinsic MHC-II expression depend on the local cellular ecosystem, the responding CD4 T cell states, and the balance between stimulatory and inhibitory cues (Johnson et al., 2018; Lei et al., 2023). Our findings posit that, in orthotopic MSS CRC under a microbiome condition that supports immune activation, the dominant effect of cancer-cell MHC-II expression is immunostimulatory rather than tolerogenic.

Considered together with the companion study by Saad et al., our findings support a model in which metastasis outcome in CRC is shaped early by the quality of local immune engagement rather than arising only from late-stage immune collapse (Saad et al., 2026; Westcott et al., 2021). Saad et al. identify divergent early T cell states during orthotopic CRC progression, whereas our study identifies a microbiome-dependent epithelial mechanism that can bias that early immune window toward productive tumor control. As such, these two studies are mechanistically complementary: one defines the T cell trajectories associated with metastatic competence, whereas the other identifies a microbiome-tumor-cell antigen-presentation axis that can promote those trajectories toward immune surveillance.

One major unresolved question is the identity of the peptides presented by tumor-cell MHC-II in this setting. This issue becomes especially interesting in light of work suggesting that antitumor CD4+ T cell responses can intersect with microbial antigens, bacterial peptide presentation, and MHC-II neoantigen biology (Alspach et al., 2019; Kalaora et al., 2021; Saad et al., 2026). It remains unclear whether cancer-cell MHC-II in CRC presents bona fide tumor antigens, mimetic antigens with structural overlap to microbial epitopes, or a mixed ligand repertoire that allows tumor immunity to intersect with the preexisting mucosal CD4+ T cell compartment. Resolving this question will require direct MHC-II immunopeptidomics, paired TCR reconstruction, and functional validation of candidate ligands across primary tumors, draining lymph nodes, and metastatic lesions.

Another open question concerns which lymphocyte populations are most important downstream of the microbiome-MHC-II axis. Our depletion and host-genetics experiments indicate that both T cells and B cells contribute to the full protective phenotype, and our single-cell analyses point to enhanced chemotactic and effector programs in both CD4+ and CD8+ compartments together with expanded B-lineage states in protective microbiome conditions. These observations are consistent with prior work showing that microbiota-specific T follicular helper cells and tertiary lymphoid structures can support antitumor immunity in CRC, but they also raise the possibility that microbe-induced expression of MHC-II on tumor cells may reinforce several distinct CD4+ programs, including Th1-like helper, cytotoxic CD4+, and B-cell-supporting states (Cui et al., 2021; Overacre-Delgoffe et al., 2021; Speiser et al., 2023). Because Saad et al. found that effective early control is associated with Th1/cytotoxic differentiation whereas pro-metastatic tumors skew toward naive-like, Tfh-like, Th17, and regulatory states, it will be important to define whether tumor-cell MHC-II selectively reinforces protective CD4+ states or more broadly increases the probability of productive local engagement (Saad et al., 2026; Westcott et al., 2021).

Our study also has translational implications. Tumor-cell MHC-II may represent both a biomarker and a therapeutic target in MSS CRC. This is consistent with prior studies suggesting that tumor-intrinsic MHC-II associate with enhanced anti-tumor immunity (Cho et al., 2021; Chung et al., 2026; Johnson et al., 2016; Rodig et al., 2018; Roemer et al., 2018). Our data extend this concept by arguing that enforced tumor-cell MHC-II can actively increase MSS CRC sensitivity to PD-1/CTLA-4 blockade. This raises the possibility that strategies that induce or stabilize tumor-cell MHC-II, whether through microbial, cytokine, or epigenetic mechanisms, could be used to convert immune-refractory MSS CRC into a more immunologically responsive state.

The Helicobacter-containing microbiome in our study should be viewed primarily as a model system for mechanism discovery rather than as a direct therapeutic target. In our experiments, single Helicobacter isolates reproduced part of the tumor-control phenotype, but suppression of metastasis was strongest with the broader transferred community, indicating that cooperative community effects matter. This point is particularly important because members of the genus Helicobacter can have divergent effects across mouse models, ranging from antitumor immune stimulation in CRC models to chronic inflammation and tumor-promoting activity in other contexts (Fox et al., 2011; Ge et al., 2019; Overacre-Delgoffe et al., 2021), Future work should therefore identify human microbes, microbial consortia, or microbial products that reproducibly induce tumor MHC-II and promote antitumor immunity.

In summary, our findings support a model in which immune surveillance of MSS CRC is shaped by whether tumor cells and the mucosal immune system enter into a productive dialogue that is facilitated by the microbiome. By identifying microbiome-dependent induction of tumor-cell MHC-II as a determinant of immune engagement, metastasis control, and response to checkpoint blockade immunotherapy, this study provides a framework for therapeutic strategies aimed at restoring epithelial antigen presentation in immunotherapy resistant MSS CRC.

## RESOURCE AVAILABILITY

### Lead contact

Further information and requests for resources or reagents should be directed to the lead contact, Semir Beyaz.

### Materials availability

All unique/stable reagents generated in this study are available from the lead contact with a completed materials transfer agreement.

### Data and code availability

- Single-cell RNA-seq data have been deposited at GEO and are publicly available under the accession numbers GSE314209 (reviewer token: gdkrsgmstpsxpmt).
- Any additional information required to reanalyze the data reported in this paper is available upon request from the lead author.

## ACKNOWLEDGMENTS

We would like to acknowledge the members of the Beyaz lab for critical discussions. We thank Cold Spring Harbor Laboratory Cancer Center Shared Resources (Animal, Flow Cytometry, Microscopy, Single Cell Sequencing, Sequencing, Organoid and Histology Core Facilities) supported in part by the National Cancer Institute Cancer Center Support Grant 5P30CA045508. The sample processing and analysis of the sequencing datasets from this study was performed on High Performance Computing (HPC) resources Elzar provided by Cold Spring Harbor Laboratory. We thank Northwell Health Biospecimen Repository and Soma Prum for facilitating acquisition of patient tissue specimens. This work was financially supported by grants to S.B from the National Cancer Institute (R37CA292807), Oliver S. and Jennie R. Donaldson Charitable Trust, the G. Harold and Leila Y. Mathers Charitable Foundation, the Mark Foundation for Cancer Research (20-028-EDV), Chan Zuckerberg Initiative/Silicon Valley Community Foundation (2021-239862), STARR Cancer Consortium (I13-0052), the Cold Spring Harbor Laboratory and Northwell Health Affiliation, New York Genome Center Polyethnic-1000 Initiative, Swim Across America.

## AUTHOR CONTRIBUTIONS

Conceptualization, S.B.; C.C.;

Methodology, C.C., J.Z., S.K.L, L.G., J.H;

Investigation, C.C., J.Z.;

Data curation, E.O.;

Formal analysis, C.C., E.O., J.Z.;

Validation, O.E., K.O., S.S.;

Writing—original draft, C.C.

Writing—review & editing, C.C., M.R., S.B.;

Funding acquisition, S.B.;

Resources, D.R., G.D., T.H., Z.S., Z.A., J.G.F., Y.Q., P.M.K.W., M.R.;

Supervision, S.B.

## DECLARATION OF INTERESTS

S.B. received research funding from Caper Labs, Elstar Therapeutics, and Revitope Oncology for research unrelated to this study.

## METHOD

## KEY RESOURCES TABLE

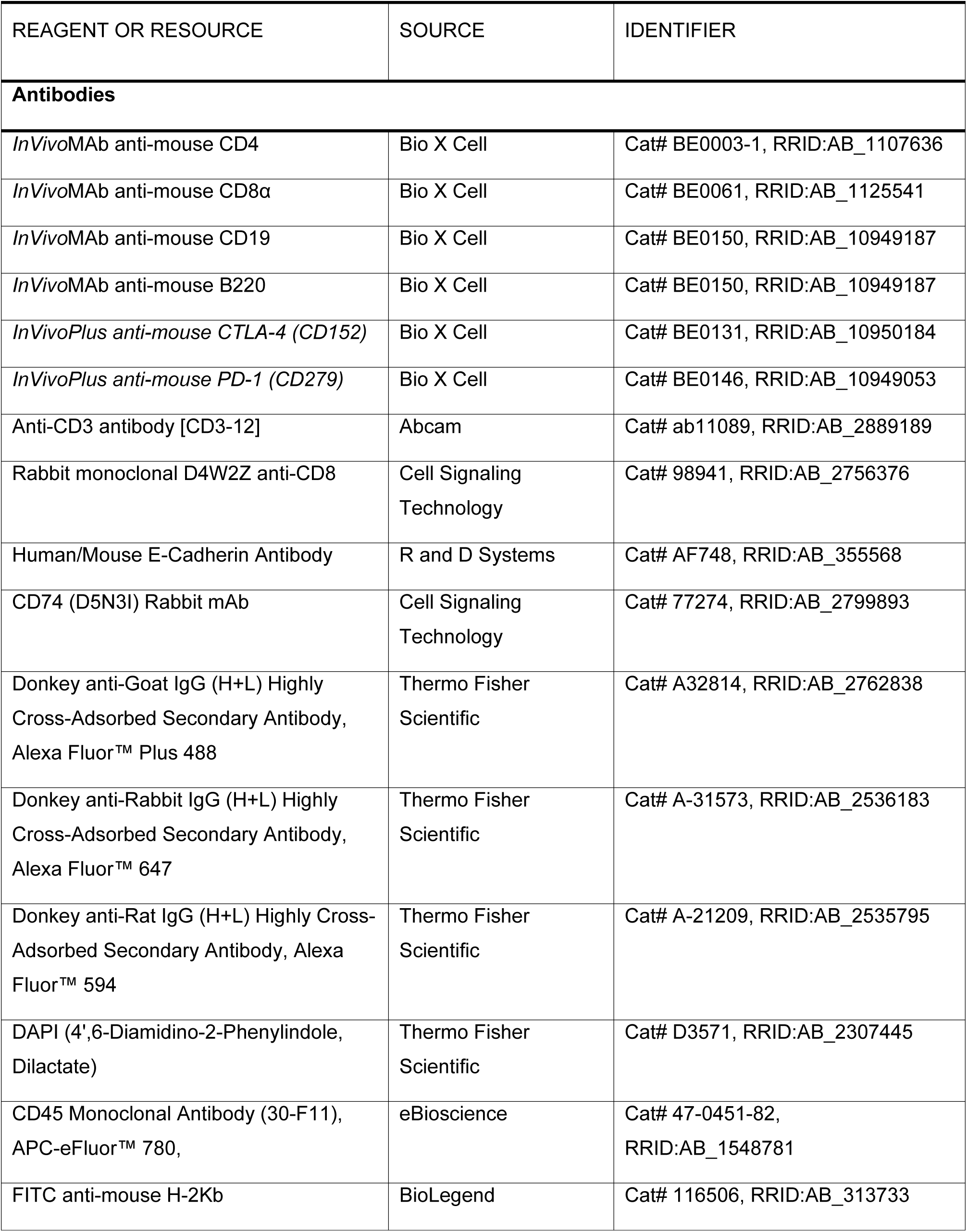
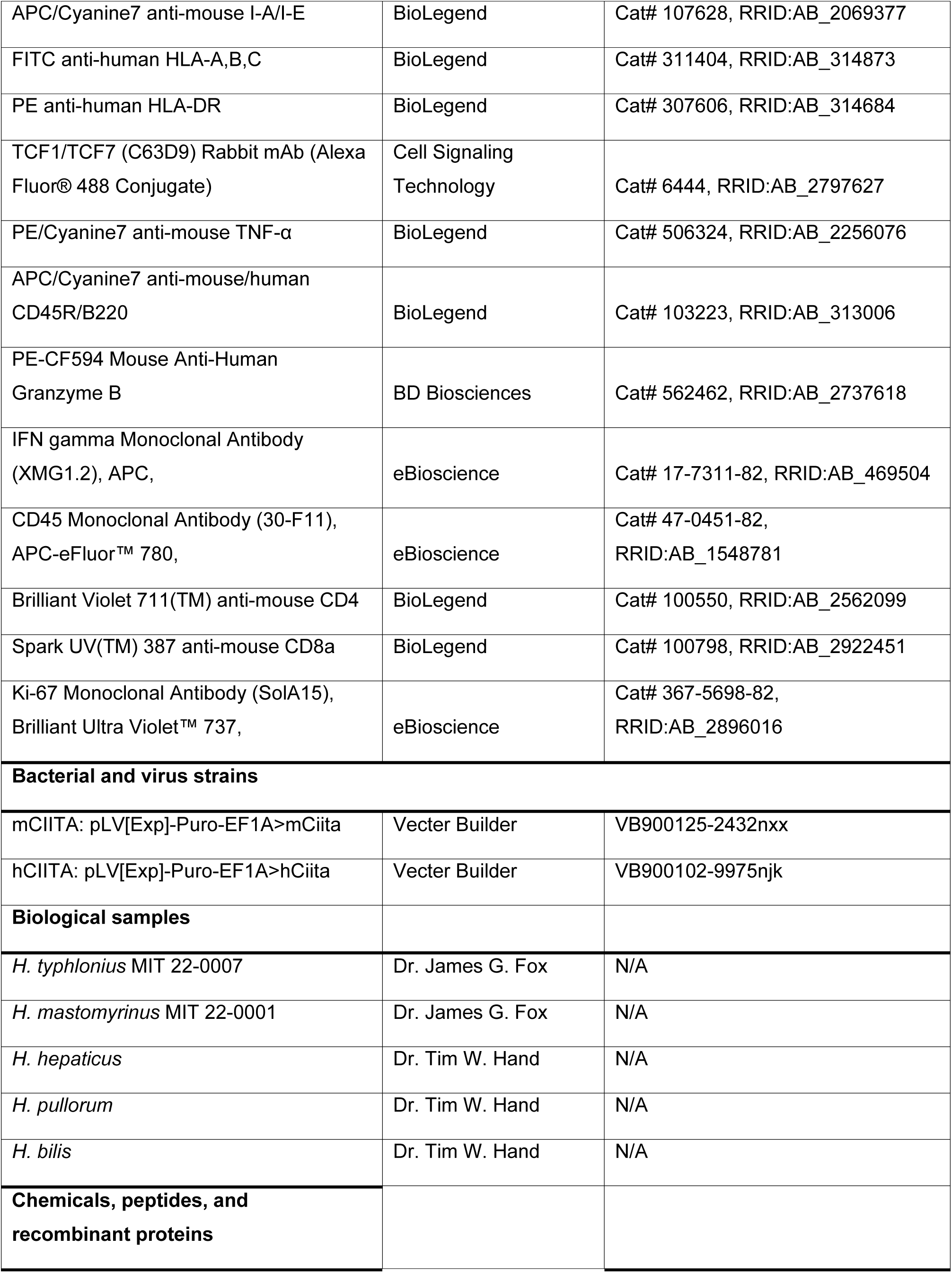
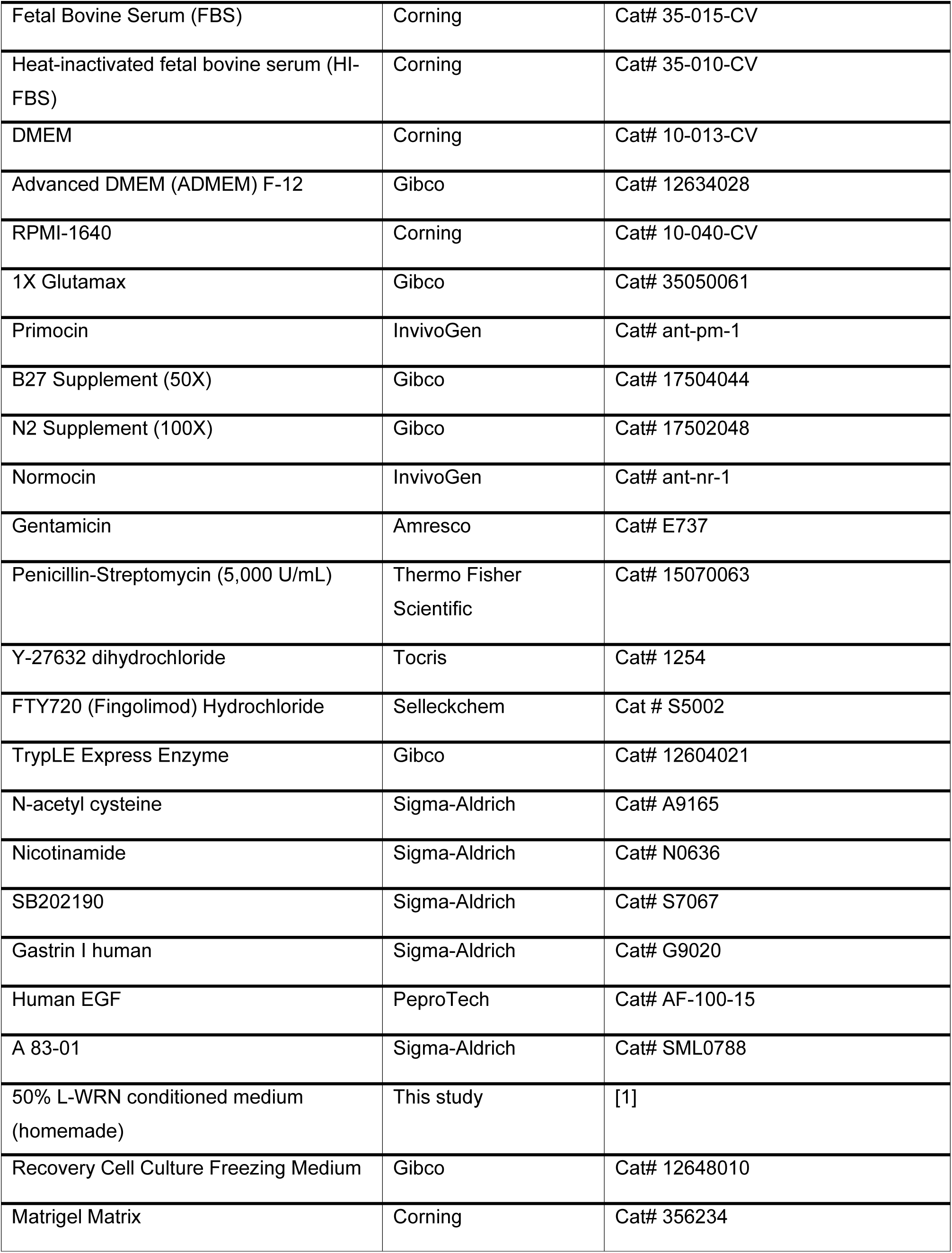
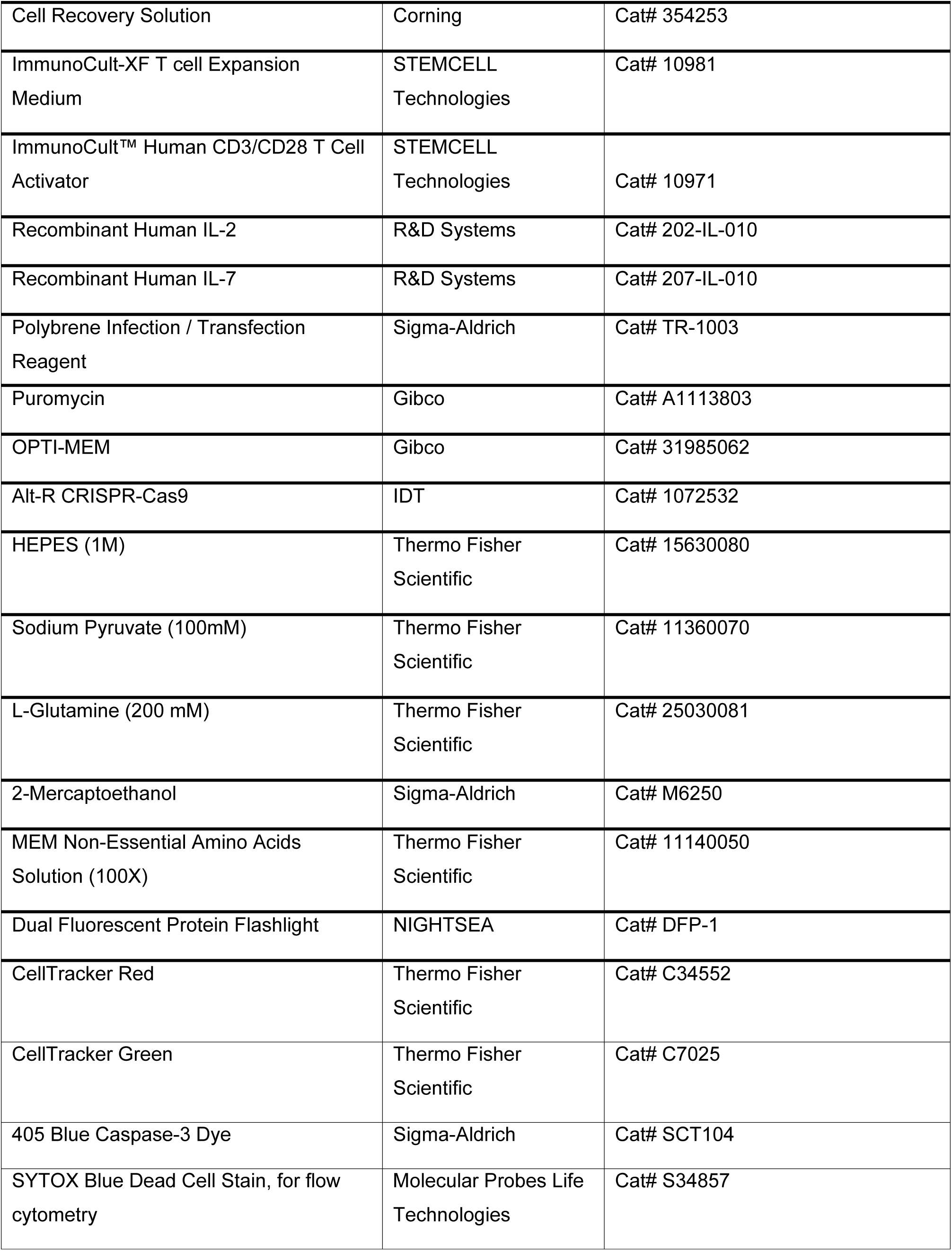
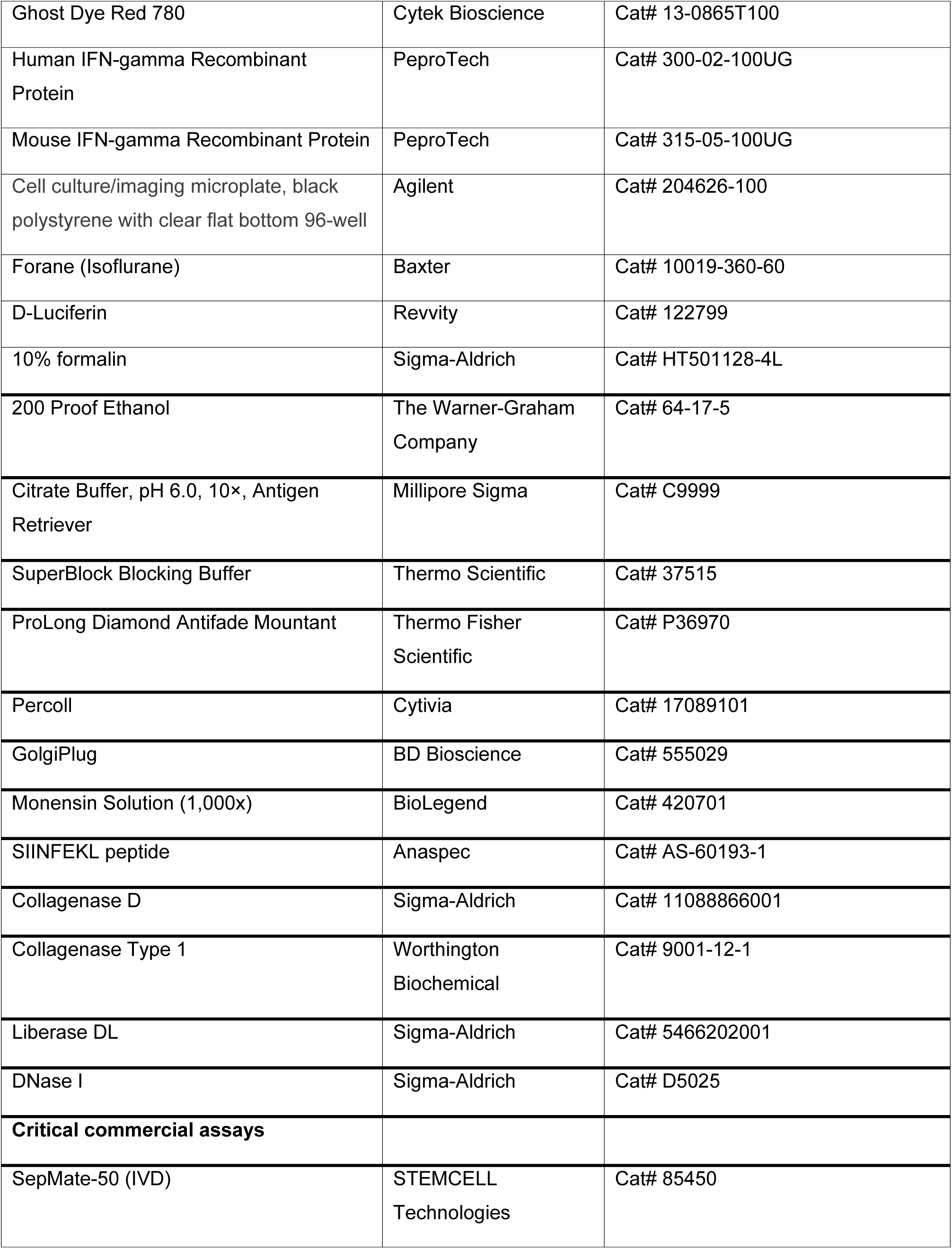
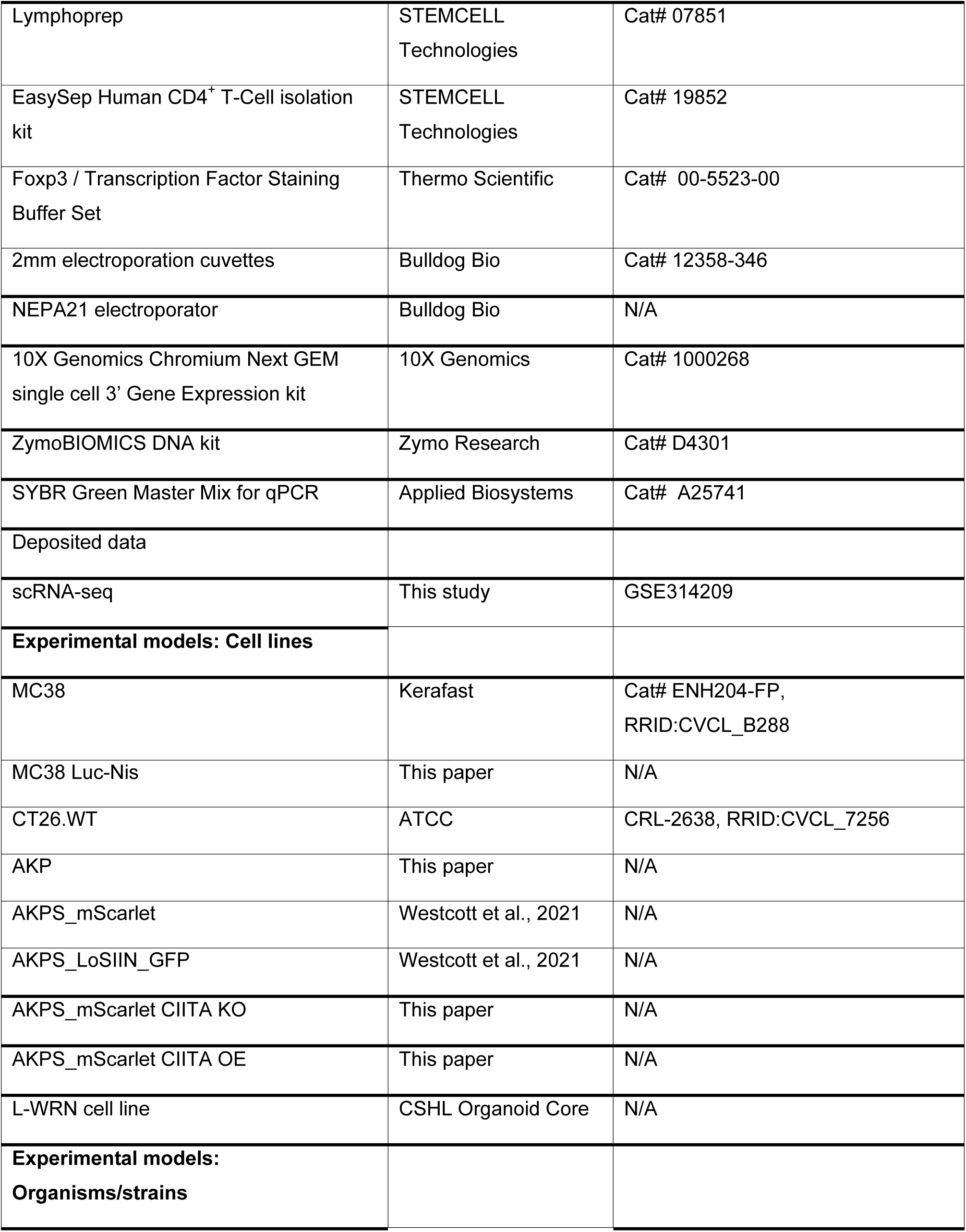
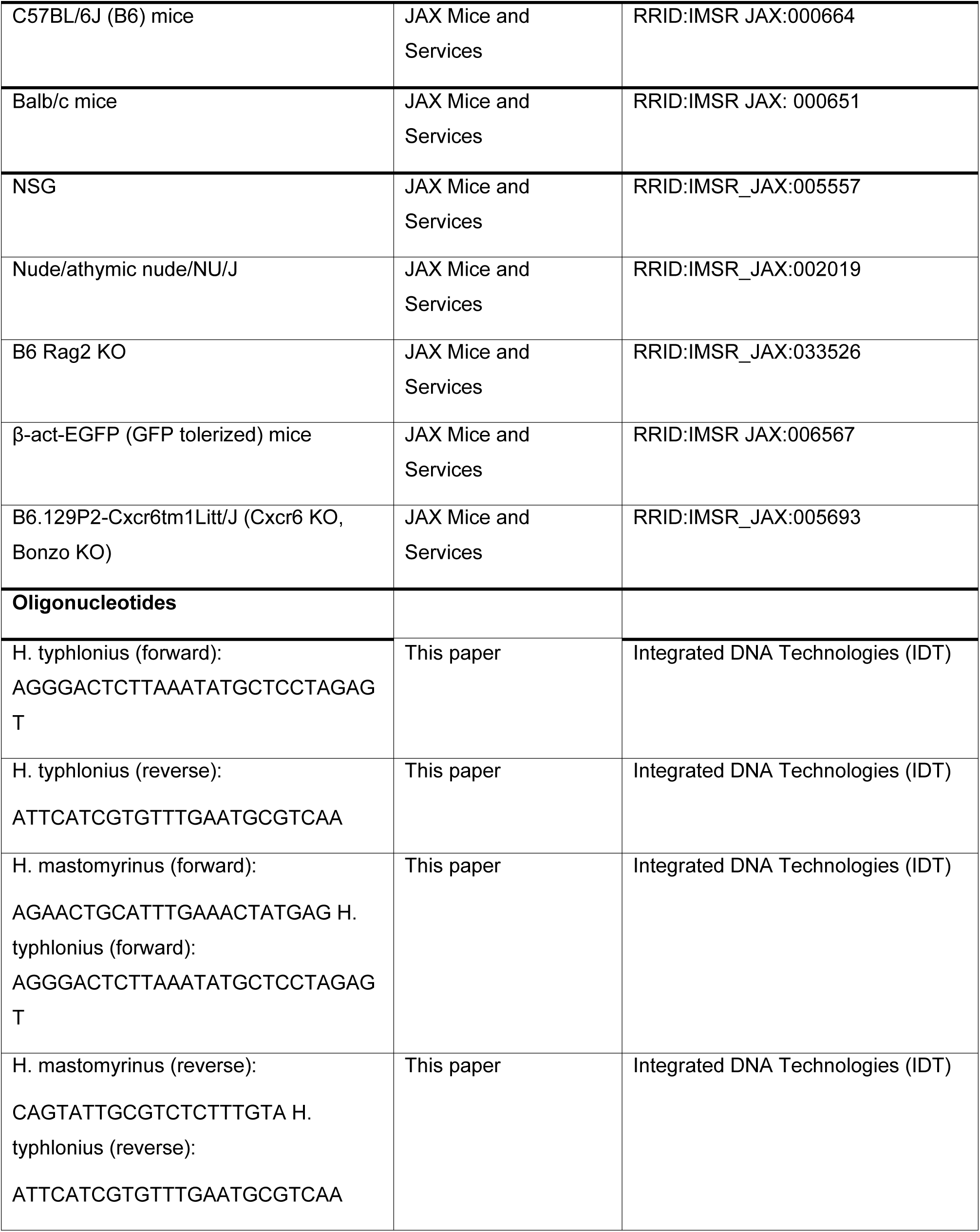
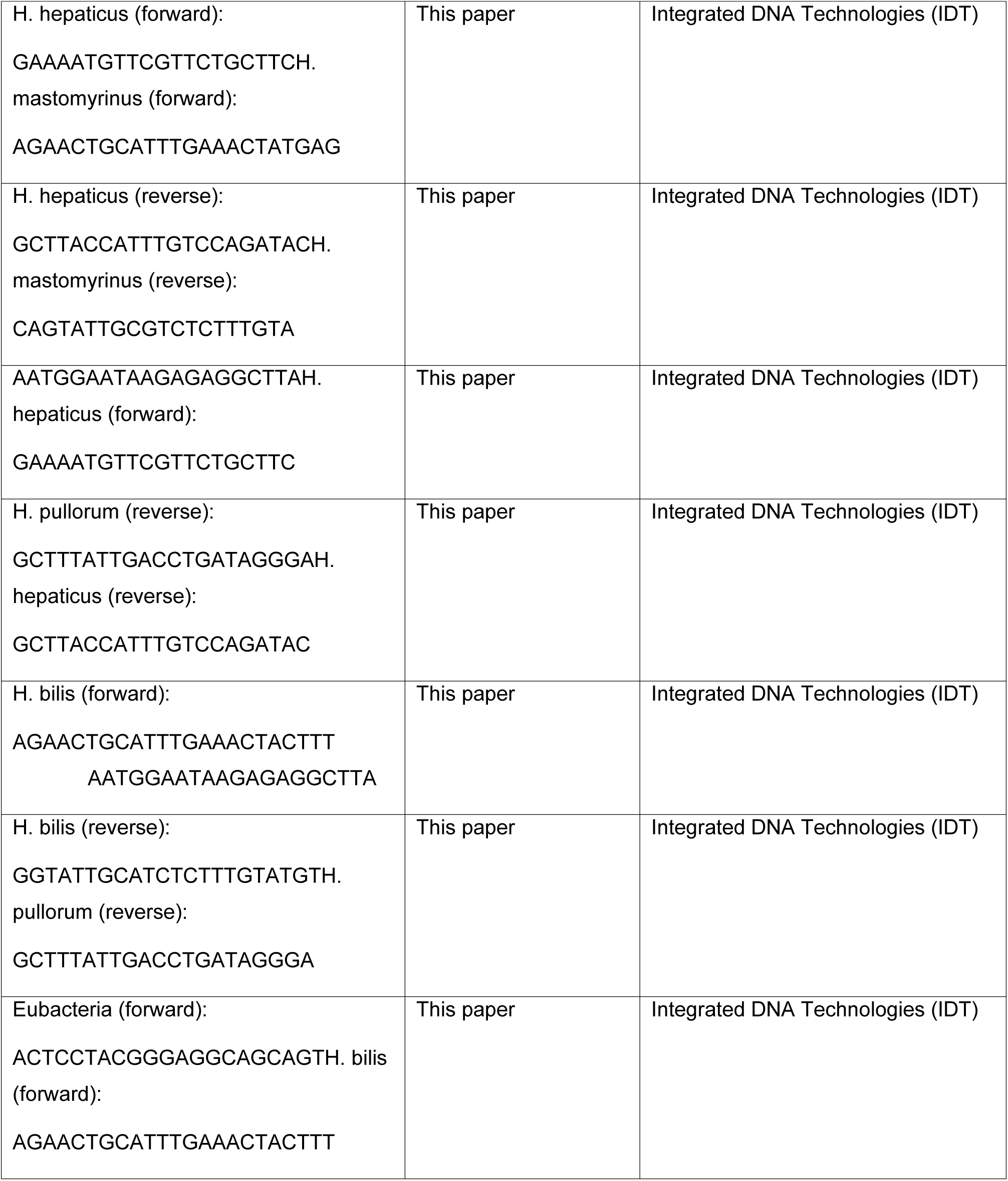
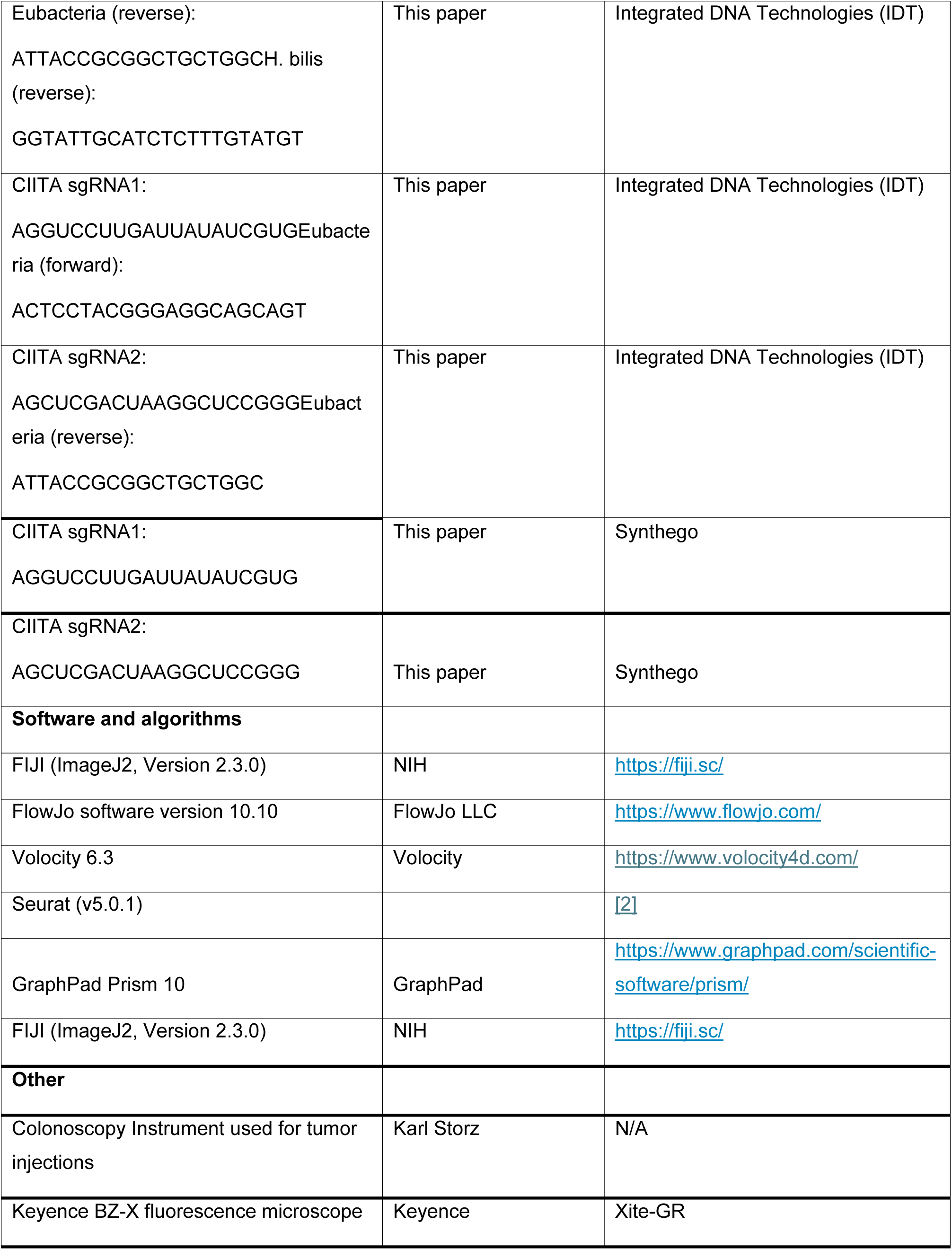
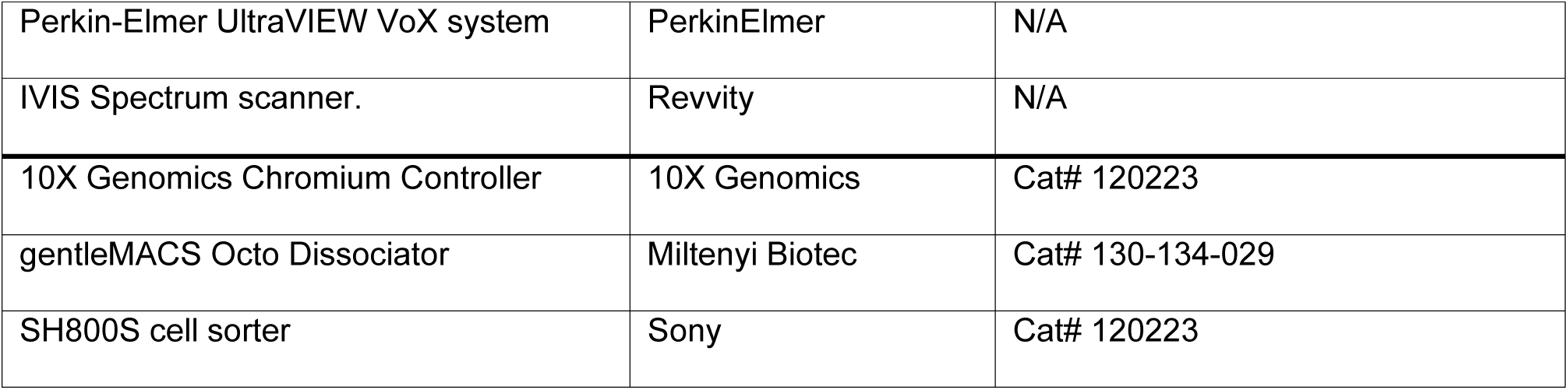

## EXPERIMENTAL MODEL AND STUDY PARTICIPANT DETAILS

### Animal models

All mouse studies were conducted in accordance with guidelines from Cold Spring Harbor Laboratory Institutional Animal Care and Use Committee (IACUC) at AAALAC accredited institution. C57BL/6J (B6) mice, Balb/c mice, B6.129P2-Cxcr6tm1Litt/J (Cxcr6 knock out) mice, NSG mice, B6 Rag2 KO mice, Nude mice, and β-act-EGFP (GFP tolerized) mice were purchased from The Jackson Laboratory. All experiments were performed with 8-10 week-old male and female mice.

### Cell lines

MC38 (Kerafast; ENH204-FP) and MC38-luc-NIS cells were maintained in DMEM (Corning) supplemented with 10% Fetal Bovine Serum (FBS), 1X Glutamax (Gibco) and Primocin (Invivogen). CT26 (ATCC; CRL-2638) cells were cultured in RPMI-1640 (Corning) supplemented with 10% FBS, 1X Glutamax, and Primocin. All cells were cultured at 37°C in humidified 5% CO_2_ incubator and passaged every 2-3 days. All cell lines were regularly tested negative for mycoplasma contamination. Research Resource Identifiers (RRIDs) for the cell lines used in this study are listed in the key resources table.

### Organoid culture

AKP (APC^KO^, Kras^G12D^, Tp53^KO^), AKPS (APC^KD^, Kras^G12D^, Tp53^KO^, Smad4^KO^ [3]), and AKPS derived organoids (AKPS loSIIN, AKPS CIITA OE, AKPS CIITA KO) organoids were embedded in Matrigel (Corning) and cultured in Minimal media (Advanced DMEM (ADMEM) F-12 (Gibco) supplemented with 1X N2 Supplement (Gibco), 1X B27 Supplement (Gibco), 1X Glutamax, and Primocin (100 μg/ml) as described previously [4]. Organoids were passaged every 4-5 days using TrypLE Express Enzyme (Gibco). AKP and AKPS organoids were fluorescently labeled, with AKP expressing GFP and AKPS expressing mScarlet enabling tumor tracing and fluorescent imaging.

### Human organoid derivation and culture

Human colon tumor samples were obtained from patients with informed consent undergoing surgical resection procedures at Huntington Hospital. Study protocols were reviewed and approved by the Northwell Health Biospecimen Repository (#1810). Tissue samples were first rinsed with cold PBS and incubated with antibiotic mixture [Normocin (100 ug/ml), Gentamicin (50 µg/ml), and 1X Pen/Strep (100U/ml Pen and 100 µg/ml Strep) for 20min on ice. Next, tissue pieces were washed again with cold PBS and digested using a digestion cocktail consisting of Collagenase (1mg/ml) and Y27632 dihydrochloride (10 µM) in DMEM, followed by incubation at 37°C for 45-60min. Upon completion of digestion, the mixture was centrifuged at 300g for 5min, and the pellets were resuspended in TrypLE express enzyme for further digest the sample into single cells for 5-10min at 37°C. Cells were then pelleted again and resuspended with 70:30 Matrigel:culture medium mixture and plated onto 6-well plates. Cells were overlaid with culture medium consisting of ADMEM, 1X Glutamax, HEPES (10mM), 50% L-WRN conditioned medium [1]), 1X B27 Supplement, 1X N2 Supplement, Nicotinamide (10mM), N-acetyl cysteine (1mM), Primocin (100 ug/ml), SB202190 (10µM), Y-27632 (10µM),Gastin I (10nM), EGF (50ng/ml), and A83-01 (500nM). Culture medium was refreshed every 4-5 days, and patient derived organoids (PDOs) were passaged every 7-10 days, typically at a 1:3 ratio.

### Human Peripheral blood mononuclear cell (PBMC) isolation

Alongside patient tissue samples, matched peripheral blood was obtained from the same individuals. Whole blood was first diluted in 1:1 with PBS containing 10% FBS, and PBMCs were isolated using SepMate-50 tubes (STEMCELL Technologies) in combination with Lymphoprep (STEMCELL Technologies). Briefly, Lymphoprep was added to the bottom of the StepMate tubes to establish the density gradient, and diluted blood was carefully layered on top. Tubes were centrifuged 1200 x g for 10min with the brake on, according to the manufacturer’s instructions. The middle layer (PBMCs) was carefully extracted using serological pipette, centrifuged and washed with PBS. Isolated PBMCs were cryopreserved in commercial freezing medium (Gibco). For co-culture assays, frozen vial was thawed and PBMCs were recovered and cultured in human T cell medium consisting of ImmunoCult-XF T cell Expansion Medium (STEMCELL Technologies) supplemented with hIL-2 (2ng/ml), hIL-7 (20ng/ml), and Primocin (100 μg/ml) until usage.

### Human T cell culture and expansion

Isolated patient PBMCs were expanded using CD3/CD28 antibody according to the manufacturer’s protocol (STEMCELL Technologies) prior to CD4^+^ T cell isolation and co-culture assays. Briefly, the 25 µl of CD3/CD28 antibody was added to 1×10^6^ PBMC in 1 mL of T cell media and cultured for 3 days. Cells were then washed, cultured with T cell media for expansion for additional 7 days. CD4^+^ T Cells were subsequently isolated using EasySep Human CD4^+^ T-Cell isolation kit (STEMCELL Technologies). The expanded immune cells, including CD4^+^ T cells, were then used for downstream co-culture assays.

### Human autologous immune cell-CRC organoids co-culture

Stable CRC organoids were split as described above and allowed to recover and grow for 2-3 days. Organoids were labeled with CellTracker Red (1µM) and autologous immune cells were labeled with CellTracker Green (1µM) for 30min, followed by two washes in their respective media. Immune cells (2 x 10^5^) were mixed with 50-100 organoids in a medium consisting of 45% organoid media, 45% human T Cell media, and 10% Matrigel supplemented with Caspase-3/7 dye (500nM). Co-cultures were plated in 96-imaging well plates (Agilent). Plates were briefly centrifuged (100 x g, 1min) to bring the cells into a single focal plane. Co-cultures were imaged following day unless otherwise stated to assess tumor-immune cell interactions and organoid apoptosis as described below.

### Live Cell Imaging and analysis

All imaging was performed using a PerkinElmer UltraVIEW VoX system, with acquisition times indicated in corresponding figures and figure legends. The system is equipped with a high-speed spinning disk (Yokogawa® CSU-X1) laser confocal microscope optimized for live cell imaging, a photokinesis unit (for FRAP and photoactivation), six laser lines (405, 440, 488, 514, 561 and 640 nm), a high-end CCD camera, and a fully automated stage enabling image tiling. Image analysis was performed using Volocity version 6.3. Z-stack images were first processed as maximum intensity projections (MIP) for downstream analysis. For each fluorophore, thresholds were set to detect positive signal from corresponding targets (e.g., Patient-derived organoids (PDOs), immune cells, and caspase-3/7) by comparison with paired brightfield images. Co-localization coefficient values [5] - defined as the ratio of a co-localized fluorophore over the total signal of that fluorophore - were calculated for individual time points and exported to Microsoft excel for further analysis. To quantify cancer-immune cell interactions, we calculated the immune cell co-localization coefficient (Immune_co-loc_ / Immune_total_) at each time point amd normalized these values to the coefficient at the initial time point (t = 0), when PDOs and immune cells were evenly distributed throughout the wells. This normalization minimized false positive values from non-interacting cells (i.e., when PDOs and immune cells were simply layered on top each other). The normalized values were plotted over the time course of co-culture. For Caspase-3/7 analyses, we calculated the co-localization coefficient of PDOs with caspas- 3/7 (Organoid_co-loc_ / Organoid_total_) and first normalized these values to the corresponding time-matched no-immune-cell control group, to account for non-immune cell control group, to account for non-immune mediated caspase-3/7 activity (e.g., apoptosis induced by starvation). The ratio between complete and incomplete apoptosis was then multiplied by these normalized coefficient values to quantify the magnitude of immune-mediated organoid apoptosis at the terminal time point.

## METHOD DETAILS

### Helicobacter species transfer

Fecal microbe transplantation was performed as described previously [6]. Briefly, two fecal pellets (∼ 50 mg total) from Helicobacter-positive donor mice were resuspended in 1 ml sterile PBS, thoroughly homogenized, and passed through a 70-μm cell strainer. Recipient mice received 200 μl of the resulting filtrate by oral gavage every other day for a total of three doses. For single *Helicobacter* species transfers (*H. hepaticus, H. pullorum, and H. bilis*), cecal contents from germ-free mice mono-colonized with the respective species were processed and administered as described above. For transfers with *H. typhlonius* or *H. mastomyrinus*, 100ul of helicobacter culture (OD=1.5) was administered by oral gavage every other day for three doses. Cecal contents were generously provided by Dr. Tim W. Hand, and single helicobacter cultures were provided by Dr. James G. Fox. Successful colonization and species identity were confirmed by quantitative PCR (qPCR) using species-specific 16S rRNA primers (Supplementary table 1) as listed in the key resources table.

### Confirmation of helicobacter colonization by fecal content qPCR

Fecal microbial DNA was extracted using ZymoBIOMICS DNA Kit (Zymo Research) according to the manufacturer’s protocol. Extracted DNA from Helicobacter-positive (whole-flora and/or single-species inoculated) and Helicobacter-negative mice was used to detect individual helicobacter species. Briefly, 10ng of extracted DNA was used per reaction for a SYBR-green (Applied Biosystems) based qPCR assay with 300nM final concentrations of species specific forward and reverse primers, as described previously [7, 8]. Cycle threshold (Ct) values were used to calculate the relative abundance of each Helicobacter species normalized to total eubacteria [8]. All primer sequences are listed in the key resources table.

### Bioluminescence imaging (BLI)

Bioluminescence imaging was performed using an IVIS Spectrum scanner (Revvity). Tumor bearing mice were first anesthetized with isoflurane (∼2% by inhalation) in an induction chamber. A filter-sterile solution of D-Luciferin (15 mg/mL in PBS) was administered by intraperitoneal (IP) injection at a dose of 10 µl per gram body weight (e.g. 200 µl for a 20 g mouse). Following injection, mice were placed on their backs in the IVIS Spectrum scanner, and anesthesia was maintained with Isoflurane throughout image acquisition. Bioluminescence images were acquired using Living Image software (Revvity). To ensure optimal substrate uptake and peak photon emission, imaging was initiated 14 minutes after IP injection. Initial scans were acquired using a 1-minute exposure time, large binning, and f-stop 1 at field of view C. At later time points, exposure time and binning were reduced as needed to prevent pixel saturation. Total anesthesia time did not exceed 30 minutes per session. Mice were imaged every two weeks to monitor tumor growth. Signal intensity was quantified as radiance (photons/sec/cm^2^/steradian) by defining a region of interest (ROI) using Living Image software.

### Cytokine treatment of organoids

Stable AKPS organoids or human CRC PDOs were split at day 4-5 of culture. Immediately after passaging, organoids were treated with either mouse Interferon-gamma (mIFNγ 2 ng/mL; PeproTech) or human Interferon-gamma (hIFNγ 2 ng/mL; PeproTech) and maintained under cytokine stimulation for 48 hours. Organoids were subsequently harvested for downstream flow cytometry assay.

### Flow cytometry

Following the 48-hour cytokine treatment, organoids (AKPS or human CRC) were digested into single-cell suspensions using TrypLE Express Enzyme. Cells were stained with an antibody cocktail containing either mouse MHC-I FITC and mouse MHC-II APC/Cy7 or human MHC-I FITC and human MHC-II PE, each at 1:200 dilution. Dead cells were excluded using the viability dye SYTOX blue (Life Technologies) from the analysis.

### Colonoscopy-guided orthotopic injection model

Orthotopic implantation of cells or organoids into the colon was performed as described previously [4, 9]. Briefly, Organoids embedded in Matrigel were dissolved with Cell Recovery Solution (Corning) at 4°C for 30 - 60 min and washed with PBS. A small portion of each organoid was digested with TripLE Express Enzyme to estimate cell number. Washed organoid suspensions were adjusted to 1×10^7^ cell-equivalent /mL for AKP organoids and 5×10^6^ cell-equivalent/mL for AKPS organoids in 10% Matrigel for injection. Intact organoids were implanted into colon sub-mucosa using a Hamilton syringe (model 7656-1) fitted with a custom 33G needle (Hamilton; custom made similar to 7803-05, 16’’ pt4’ Deg12). Each mouse received approximately 100ul of adjusted cell suspension (1×10^6^ cell-equivalent for AKP and 5×10^5^ cell-equivalent for AKPS organoids per injection). Successful injections were confirmed by appearance a visible mucosal bleb in the colon. Orthotopic injections of 2D cell lines (MC38, MC38-luc-NIS, and CT26) were performed similarly, using 1×10^7^ cells/mL for MC38 and MC38-luc-NIS and 2.5×10^6^ cells/mL for CT26. Tumor growth was monitored by optical colonoscopy using Karl Storz Image 1 HD Camera System (Image 1 HUB CCU, 175 Watt Xenon Light Source) equipped with a Richard Wolf 1.9mm/9.5 Fr integrated telescope (Richard Wolf, model 8626.431). Mice were euthanized at specified time points or upon reaching humane endpoints (either loss of 20% body weight or displaying poor body condition, i.e., hunched posture). Metastatic sites (liver, lung, omentum, and rarely lymph nodes) were screened for mScarlet fluorescence using a fluorescence excitation flashlight (Electron Microscopy Sciences; Xite-GR) and confirmed by histological analysis. Mice with failed injections, defined as absence of tumor formation by day 7 or isolated tumors confined to the intraperitoneal space, were excluded from analysis.

### Tumor index and size measurement

Colonoscopy images were recorded and analyzed offline. Tumor index was calculated as described previously [10]. Briefly, tumor size was measured from colonoscopy images and normalized to the diameter of the lumen, where absence of detectable tumor corresponds to a tumor index of 0 and complete luminal obstruction by tumor corresponds to a tumor index of 1. For normalized tumor index, tumor index values obtained after treatment regimens (e.g., Helicobacter inoculation or immunotherapy) were normalized to corresponding tumor index values measured prior to the initiation of the treatment regimen.

For gross tumor size measurement, tumors were harvested at day 21 by dissecting the tumor-bearing region of the distal colon. Harvested tumors were then lateralized and trimmed to exclude adjacent non-mScarlet-expressing tissue. Images were acquired with a ruler placed next to each specimen to measure tumor length and width, while height was measured separately with a ruler. Tumor volume was calculated by multiplying the measured length x width x height for each specimen.

### *In vivo* FTY720 treatment

For *in vivo* administration of FTY720 (Fingolimod; Selleckchem; S5002) was dissolved in DMSO and diluted to a working concentration of 5µg/ml. Mice were treated with FTY720 (1mg/kg) by daily oral gavage starting one day prior to tumor implantation and continuing for 14 days. Tumors were collected on day 15.

### Immunotherapy of tumor bearing mice

AKPS organoids (5×10^6^ cell-equivalent) were orthotopically implanted into the colon of recipient mice as described above, and successful tumor establishment was confirmed by colonoscopy at day 7 post-implantation. Beginning on day 21 post-implantation, mice were treated intraperitoneal injection with anti-PD1 antibody (200µg per dose; BioXcell) and anti-CTLA4 antibody (200µg for the initial dose followed by 100µg for subsequent doses; BioXcell). Antibodies were administered every other day for two weeks for a total of six doses. Tumor progression was monitored bi-weekly by colonoscopy throughout the treatment course.

### Preparation of formalin-fixed paraffin-embedded (FFPE) tissue slides

For histological analysis, tissues (colon, liver, and lungs) were harvested from mice at the indicated experimental time points. For colon tumors, colon was opened longitudinally prior to the embedding process. Tissues were fixed in 10% formalin for 24-48 hours at room temperature and subsequently transferred to 70% ethanol for storage prior to processing. Fixed tissues were embedded in paraffin, and tissue blocks were sectioned at a thickness of 10-µm. Sections were submitted to CSHL Animal and Tissue Imaging Shared Resources for hematoxylin and eosin (H&E) staining.

### Immunofluorescence staining

FFPE tumor sections were deparaffinized in xylene and rehydrated through a graded ethanol series. Antigen retrieval was performed in citrate buffer (Millipore Sigma) using microwave. Sections were blocked with SuperBlock for 1 hour at room temperature and incubated overnight at 4°C with primary antibodies against CD3 (Abcam; 1:100), CD8a (1:200; Cell Signaling Technology), E-Cadherin (1:100; R&D system). After three washes in 0.05% PBS-T, slides were incubated with species-appropriate secondary antibodies [Goat IgG (1:200; Invitrogen), Rabbit IgG (1:200; Invitrogen), and Rat IgG (1:200; Invitrogen;)] for 1 hour at room temperature. Sections were then washed twice with PBS, counterstained with DAPI (0.5ug/ml; Invitrogen) for 5 min and mounted. Slides were imaged using a Keyence BZ-X fluorescence microscope (Keyence) and processed in FIJI (ImageJ2, Version 2.3.0). Quantification was performed manually as described below.

### Image quantification

Cancer cell-contacting CD8^+^ immune cells was quantified by using Imaris. Briefly, Epcam channel was used to generate a surface that contained all glandular tumor structures within each field-of-view. This surface was then used as spatial mask to restrict analysis in the remaining channels to regions immediately adjacent to EpCAM^+^ tumor structures. Co-localization of CD3 and CD8 signals was used to identify and quantify cancer cell-contacting CD8^+^ T cells using Imaris “Spots” detection module. For CD74 quantification, regions of interest (ROI) within tumor areas were manually drawn and mean fluorescent intensity was calculated for each ROI.

### Lentiviral transduction of organoids

To generate CIITA overexpression (CIITA OE) lines, AKPS and human colorectal cancer (CRC) organoids were digested into single cells using TripLE express Enzyme, washed, and resuspended in complete organoid culture medium. 1×10^6^ cells were incubated with 10 µl of lentiviral particles encoding either mouse CIITA (pLV[Exp]-Puro-EF1A>mCiita) or human CIITA (pLV[Exp]-Puro-EF1A>hCiita), both from Vector Builder, in the presence of polybrene (20 µg/ml; Sigma). Cells were plated into individual wells of a 12-well plate and centrifuged at 300 x g for 1h room temperature (spinfection), followed by incubated at 37°C in a humidified 5% CO_2_ incubator for an additional 3 hours. Cells were then pelleted, resuspended in a mixture of 30% medium and 70% Matrigel, and plated into a 6-well plate. After 4-5 days, organoids were passaged and subjected to puromycin selection (7µg/ml; Gibco). Surviving organoids were expanded and subsequently analyzed by flow cytometry to confirm successful CIITA overexpression.

### Organoid CRISPR/Cas9 RNP electroporation

To generate CIITA knockout (CIITA KO) AKPS organoid lines, day 4 AKPS organoids were dissociated into single cells as described above and resuspended in 100 μl OPTI-MEM (Gibco). Ribonucleoprotein (RNP) complexes were assembled by mixing 1.64 μl (0.1 nmol) Alt-R Cas9 nuclease (Integrated DNA Technologies) with 3 μl (0.3 nmol) synthetic sgRNA (Synthego) and incubating for 10–20 min at room temperature. Cells were then added to the RNP mixture, transferred to a 2-mm gap electroporation cuvette (Bulldog Bio) and electroporated using a NEPA21 electroporator (Bulldog Bio) with the following settings: poring pulse parameters: 175 V, 5-ms pulse length, 50-ms interval, 2 pulses, 10% decay rate, + polarity; and transfer pulse parameters: 20 V, 50-ms length, 50-ms interval, 5 pulses, 40% decay rate, ± polarity. Following electroporation, organoids were gently resuspended in pre-warmed minimal medium and incubated at 37 °C for 15 minutes prior embedding in Matrigel. Electroporated organoids were allowed to recover and subsequently passaged. Newly split organoids were then treated with mouse IFN-γ (2ng/ml), to induce MHC II expression, followed by sorting on a Sony SH800S cell sorter to isolate MHC-II-negative CIITA KO cells.

### Immune cell priming

#### Tissue preparation

To distinguish tissue-infiltrating from circulating T cells, mice were injected intravenously with APC-eFluor 780 CD45 antibody (eBiosciences) shortly before euthanasia. Colon-draining lymph nodes (caudal and iliac) were harvested and mechanically dissociated in RPMI-1640 (Corning) supplemented with 5% heat-inactivated fetal bovine serum (HI-FBS) (Corning). Tumors were identified using a Dual Fluorescent Protein Flashlight (Nightsea), excised and transferred into digestion buffer containing Collagenase Type 1 (500 U ml−1; Worthington) and DNase I (20 μg ml−1; Sigma-Aldrich). Tumor were finely minced with surgical scissors and digested at 37 °C for 40 min with gentle agitation. Following enzymatic digestion, tumors were further dissociated using a gentleMACS Octo Dissociator (Miltenyi Biotec) with the tumor_imp1.1proggram and passed through a 100-μm cell strainer to generate single-cell suspensions.

Lymphocytes from the cell suspension were isolated by gradient isolation using Percoll (Cytivia). Briefly, digested tissues were mixed with 5 mL of 44% Percoll in DMEM, and then 3 mL of 67% Percoll in PBS was underlaid by glass Pasteur pipette. Without disturbing the interface, samples were spun down at 2000 rpm for 20min without brake at room temperature. Upon completion of centrifugation, buffy coat was transferred into a 15 ml conical tube, topped off with RPMI 1640 media, and spun down at 2000 rpm for 10min at 4°C. Lymphocytes from colon-draining lymph nodes (dLNs) and tumor were divided for either immediate flow cytometric staining or *ex vivo* peptide stimulation.

#### Flow cytometry staining and analysis

Live/dead discrimination was performed using Ghost Dye Red 780 (1:500; Cytek Bioscience) diluted in PBS. Surface staining was carried out in FACS buffer consisting of 1 mM EDTA, 25 mM HEPES, 0.5% HI-FBS in PBS. Cells were fixed for 1 hour at room temperature using Fixation/Permeabilization Concentrate (eBioscience) diluted 1:3 in Fixation/Permeabilization diluent (eBioscience) and washed in permeabilization buffer (eBioscience). Intracellular staining was performed in permeabilization buffer overnight at 4 °C. After staining, cells were washed and resuspended in FACS buffer for acquisition on a BD LSRFortessa flow cytometer (BD Biosciences) using BD FACSDiva software v8.0. Data were analyzed using FlowJo v10.4.2 (FlowJo). Single lymphocytes were identified by FSC-A versus SSC-A gating followed by FSC-A versus FSC-H to exclude doublets. Live CD8+ T cells were defined as CD8α^+^ Ghost Dye^-^cells. Expression of additional markers was assessed specifically within this antigen-specific CD8^+^ T-cell population for all T cell-related flow cytometry experiments in this study. For CD4^+^ T cells, live CD4^+^ Ghost Dye^-^ cells were gated, and subsequent marker expression was evaluated within this population.

#### Peptide stimulation for cytokine staining

Single-cell suspensions were prepared from tumors and draining lymph nodes as described above. Immune cells were then stimulated in mouse T-cell medium consisting of RPMI-1640 supplemented with 10% HI-FBS, 20 mM HEPES, 1 mM sodium pyruvate, 2 mM l-glutamine, 50 μM β-mercaptoethanol, 1× non-essential amino acids and 0.5X (50U/ml Pen and 50 µg/ml Strep) penicillin/streptomycin. Cells were incubated with GolgiPlug (1:1000; BD Bioscience), Monensin Solution (2 μM; BioLegend) and SIINFEKL peptide (Anaspec; 1 μM; AS-60193-1) for 3 hours at 37 °C. Following stimulation, cells were washed and stained for surface and intracellular markers as described above.

### Single cell RNA sequencing

Tumors were minced and digested in digestion medium consisting of RPMI1640 (Corning) supplemented with 10% FBS, collagenase D (2.5mg/ml; Sigma-Aldrich), Liberase DL (0.5mg/ml; Sigma-Aldrich), and DNase I (0.2mg/ml; Sigma-Aldrich). Digestion was carried out at 37°C with rotation at a slow speed (25 rpm) for 30-60 minutes. Digested samples were filtered through 100-µm cell strainers, and remaining tissue fragments were mechanically disrupted using plunger of a syringe. Cells were centrifuged at 300 x g for 5 minutes, washed once with RPMI1640, and stained with CD45-APC/Cy7 (immune cells) or EpCAM-APC (epithelial cells) antibodies (1:200, BioLegend). CD45^+^ EpCAM^-^ cells and CD45^-^ EpCAM^+^ cells were sorted using Sony SH800S cell sorter and combined at an 80:20 ratio, followed by single cell library preparation.

Single cell libraries were generated using a Chromium Controller (10X Genomics) using Chromium Next GEM single cell 3’ Gene Expression kit (10X Genomics), according to manufacturer’s instructions at CSHL Single Cell Biology Shared Resource. Cell suspensions were loaded to target 8000 cells per sample. cDNA and libraries were assessed using Agilent 2100 Bioanalyzer, quantified by KAPA qPCR, and sequenced on either a NextSeq500 or Novaseq2000 (Illumina) to a depth of approximately 20.000 reads per cell. FASTQs files aligned to the mouse reference genome (gex-mm10-2020-A) and digital gene-cell count matrices were generated using default parameters.

### Single cell RNA sequencing data analysis

#### MC38

Data quality was assessed according to the published guidelines [11, 12]. Cells with >10% mitochondrial transcripts, < 200 or > 2500 detected genes, >10,000 RNA counts, or <250 RNA counts were excluded. Following quality control (QC), data was processed using Seurat (v5.0.1) [2] for normalization, clustering, and integration. Each dataset was normalized using SCTransform, and the 5,000 most variable genes were identified using SelectIntegrationFeatures. Samples were integrated using PrepSCTIntegration, FindIntegrationAnchors, and IntegrateData. Following integration, Ptprc-negative cells (Ptprc≤0) were extracted using Seurat’s FetchData function. The Ptprc-negative dataset was clustered using 15 principal components (PCs) at a resolution of 0.1, and clusters were annotated based on marker gene expression. Clusters corresponding to the non-immune compartment (clusters 1, 7, and 8) were selected and reprocessed using 10 PCs at a resolution of 0.1. Cell identifiers from this non-immune subset were then removed from the main population to generate the immune compartment, which was processed using 35 PCs at a resolution of 0.6. Low-quality clusters were removed, and the immune dataset, and cell types were annotated based on the results from FindAllMarkers and established literature. UMAP was applied using the corresponding PC dimensions for visualization. Figures were generated using DimPlot, DotPlot, and dittoBarPlot package (https://www.bioconductor.org/packages/release/bioc/html/dittoSeq.html). Gene signature scoring was calculated using UCell (https://github.com/carmonalab/UCell), with gene sets for provided in the supplementary table 2 (MC38) and 3 (AKPS). Violin plots of signature score were generated using ggplot2 (https://cran.r-project.org/web/packages/ggplot2/citation.html) to visualize differences between experimental conditions. For each signature, distributions were shown as violins scaled to the data distribution and overlaid with box plots indicating median and quartile values. Statistical significance between groups was assessed using the geom_signif function’s t-test option (https://github.com/const-ae/ggsignif). Differential gene expression between experimental conditions was evaluated using Libra’s run_de function with the Wilcox method (https://github.com/neurorestore/Libra).

#### AKPS

Data quality was assessed according to the published guidelines [11, 12]. Cells with >10% mitochondrial transcripts, < 200 or more > 5,000 detected genes, >25,000 RNA counts, or <250 RNA counts were removed. Following quality control (QC), data were processed using Seurat (v5.0.1) [2] for normalization, clustering, and integration. Six samples from two sequencing runs (CC01 and CC02) were combined and processed following the Seurat v5 integration workflow (https://satijalab.org/seurat/articles/seurat5_integration tutorial). Datasets were integrated using the IntegrateLayers function with the CCAIntegration method. Post-integration, clustering was performed using 20 principal components (PCs) at a resolution of 0.2. Based on cluster marker expression, candidate non-immune cells (clusters 6 and 8) were identified and confirmed by absence of Ptprc expression (≤0). The non-immune subset was further reprocessed using 10 PCs at a resolution of 0.1, during which low-quality cells (cluster 4) were removed. The immune fraction was then generated by removing all non-immune cell identifiers from the initial dataset and re-clustering the remaining cells using 20 PCs at a resolution of 0.2. To further resolve T cell subpopulations, T cells within the immune object were sub-clustered using 25 PCs at a resolution of 0.1. Because the T cell gene expression did not clearly separate all subsets, ambiguous clusters were further sub-clustered and annotated based on CD3, CD4, and CD8 expression. The refined T cell clustering was then incorporated back into the main T cell object. Visualization and differential gene expression analysis were performed using the same workflows described for the MC38 dataset, including UMAP visualization, gene signature scoring and statistical testing.

### *In vivo* antibody mediated immune cell depletion

For T cell depletion, mice were injected intraperitoneally with anti-CD8 (200 µg per dose; BioXCell) and anti-CD4 (200 µg per dose; BioXCell) on days −1, 1, 3, 6, 8, and 10 relative to tumor implantation. For B cell depletion, mice were injected intraperitoneally with anti-CD19 (200 µg per dose; BioXCell) and anti-B220 (200 µg per dose; BioXCell) using the same dosing schedule. Following the initial two-week depletion phase, mice received 100 µg of the respective antibody combinations of antibodies once weekly for the remainder of the experiment to maintain immune cell depletion.

### Data availability Statement

scRNA-seq data can be accessed from Gene Expression Omnibus (GEO) with the following accession number: GSE314209

## QUANTIFICATION AND STATISTICAL ANALYSIS

Statistical analysis for experimental data was performed using GraphPad Prism. Statistical significance of differences between groups was assessed using unpaired t-tests, except for survival (Kaplan-meier curve). Error bars in figures indicate standard error of the mean (SEM). Statistical details, including statistical tests used, number and nature of samples analyzed are described in the legend for each figure panel. Statistical analyses for gene expression data were conducted in R.

---

**Supplementary Figure 1.**
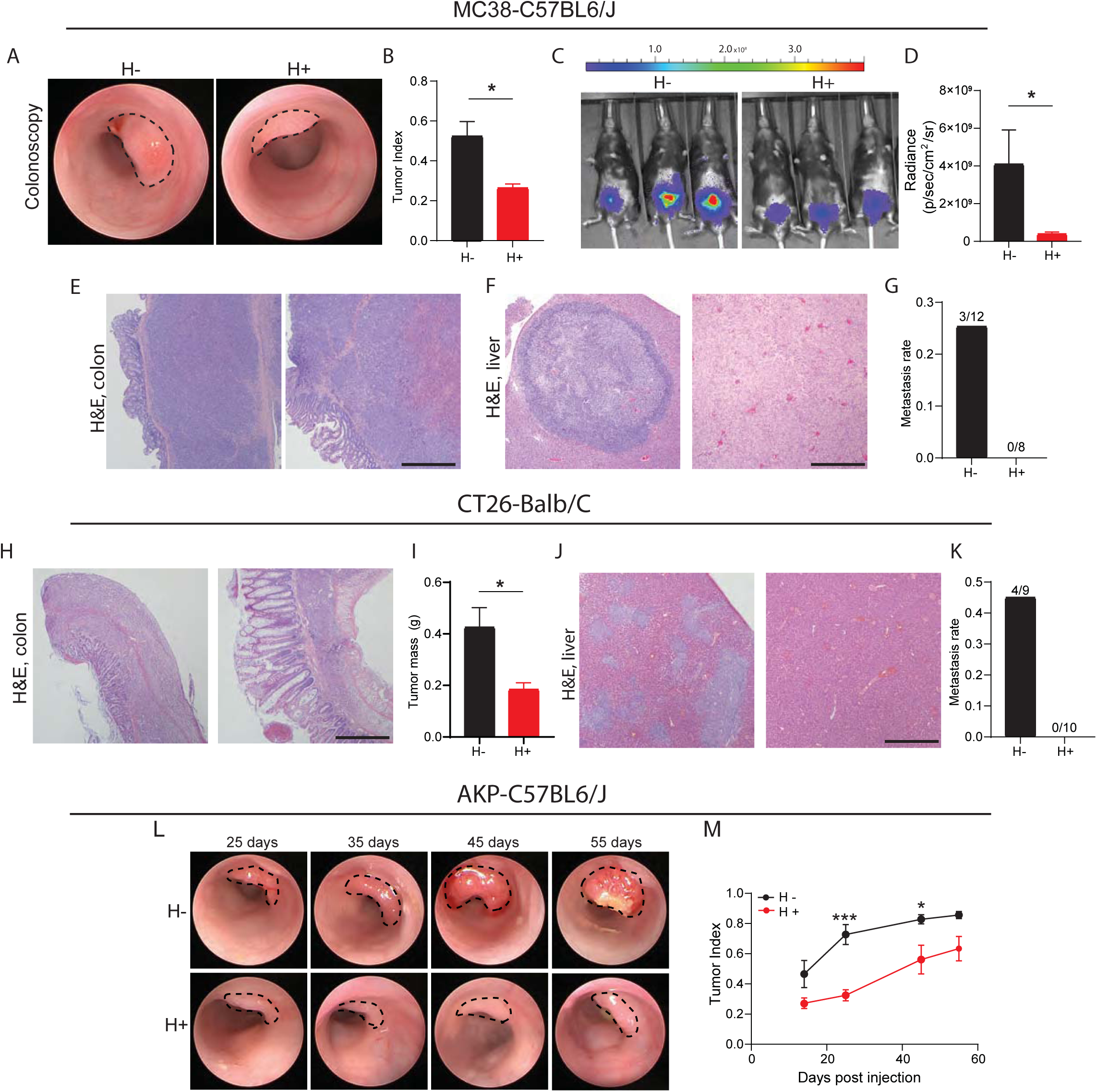
Helicobacter containing gut microbiome regulates tumor growth and metastasis across different mouse strains. (A) Representative colonoscopy images of colon tumors induced by MC38 cells in Helicobacter negative (H-) and Helicobacter positive (H+) C57BL6 mice at day 14. (B) Tumor index at day 14 after MC38 implantation (H-, n=12; H+, n=8). (C) Representative bioluminescence imaging by luciferase-labeled MC38 tumors in the colons of H- and H+ mice. D-luciferin injections (150mg/kg), imaged after 14min post injection. (D) Quantification of average photon values of tumor burden shown in (C) (H-, n=3; H+, n=5). (E - F) Representative H&E images of primary colon tumors (E) and liver metastasis (F) arising from MC38 tumors in H- and H+ C57BL6/J mice. Scale bar, 500μm. (G) Liver metastasis rate in MC38 tumor-bearing mice (H-, n=12; H+, n=8). (H) Representative H&E images of colon tumors induced by CT26 cancer cells in H- and H+ Balb/cJ mice. (I) Tumor mass of CT26-induced tumors 2 weeks after implantation in H- and H+ Balb/cJ mice (n=5, per group). (J) Representative H&E images of liver metastasis arising in CT26 tumor-bearing H- and H+ Balb/C mice. (K) Liver metastasis rate in CT26 tumor-bearing mice (H-, n=9; H+, n=10). (L) Representative timelapse colonoscopy images of colon tumors induced by AKP organoids in H- and H+ mice. (M) Longitudinal tumor index quantification depicted in (L) (H-, n=9; H+, n=7). Unpaired *t*-test and fisher’s exact test (G, K) were used for statistical analysis (B, D, I, M). (* p<0.05, *** p<0.01).

**Supplementary Figure 2.**
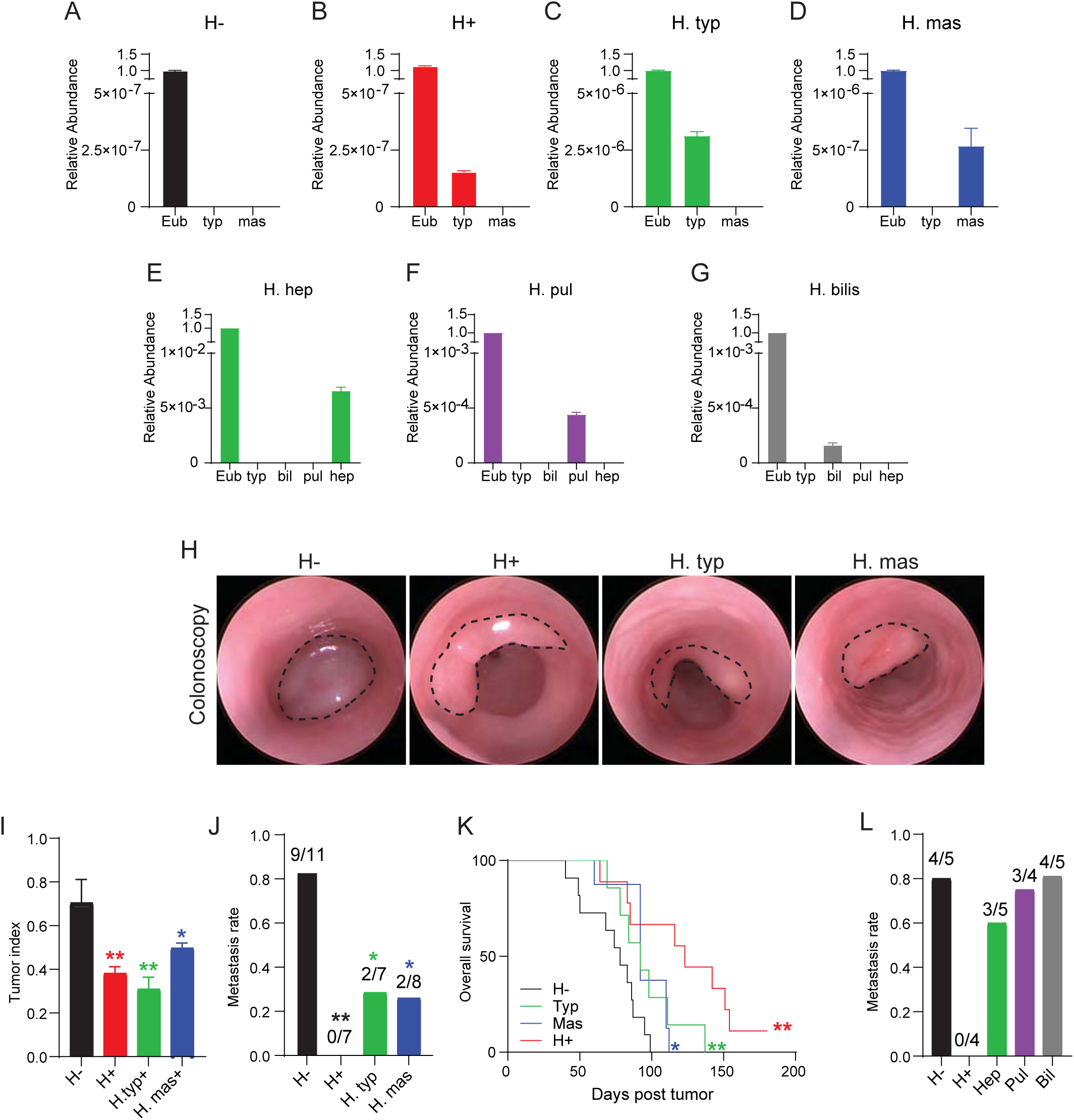
Single Helicobacter species inoculation is sufficient to elicit anti-tumor effects against MSS CRC. (A-D) Relative (to Eubacteria) abundance of H. typhlonius and H. mastomyrinus in feces from H-(A), H+ (whole flora transfer, B), and single culture inoculated (C, D) mice (n=2 each). (E-G) Relative (to Eubacteria) abundance of H. typhlonius, H. bilis, H. pullorum, and H. hepaticus in feces from H+ (whole flora transfer, E), and single culture inoculated (F, G) mice (n=4 each). (H) Representative colonoscopy images of colon tumors induced by AKPS organoids in H-, H+ (whole flora), *H. typhlonius* single culture inoculated, and *H. mastomyrinus* single culture inoculated mice at day 42 (H-, n=11; H+, n=7; H. Typ., n=7; H. Mas., n=8). (I) Tumor index 6 weeks after AKPS organoid implantation in the indicated groups (H-, n=11; H+, n=7; H. Typ., n=7; H. Mas., n=8). (J) Liver metastasis rates in the indicated groups bearing AKPS-induced tumors (H-, n=11; H+, n=7; H. Typ., n=7; H.Mas., n=8). (K) Survival rate in the indicated groups bearing AKPS-induced tumors (H-, n=11; H+, n=7; H. Typ., n=7; H. Mas., n=8). Kaplan-Meier analysis was used for survival comparisons (* p<0.05, ** p<0.01). (L) Liver metastasis rate of H-, H+ (whole flora), single-culture *Helicobacter hepaticus* (Hep), *Helicobacter pullorum* (Pul) and *Helicobacter Billis* (Bil) mice bearing AKPS-induced tumors (H-, n=5; H+, n=4; H. Hep., n=5; H. Pul., n=4; H. Bil., n=5). Ordinary one-way ANOVA was used for tumor index comparisons (B), and Fisher’s exact test was used for metastasis frequency analysis (C, E). (* p<0.05, ** p<0.01).

**Supplementary Figure 3.**
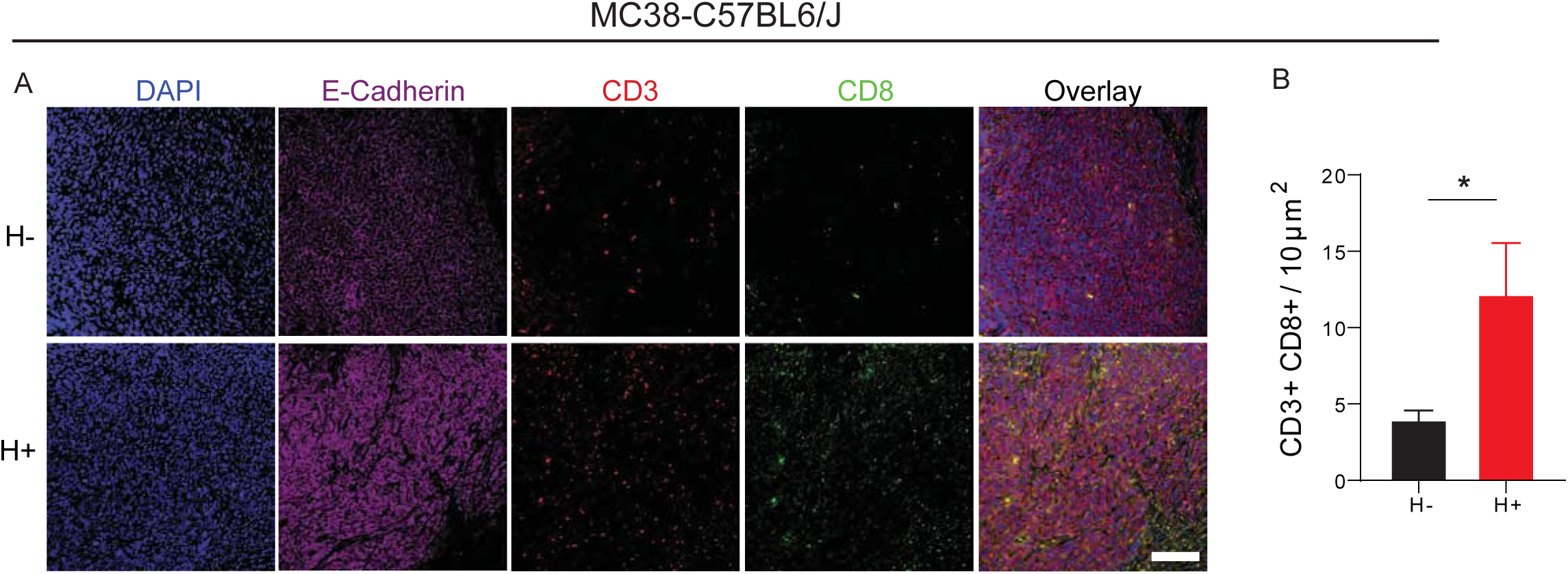
Helicobacter colonization promotes the infiltration of TILs in MC38 tumors. (A) Representative immunofluorescence (IF) images of MC38 tumors Bar=50μm. DAPI (blue), E-cadherin (white), CD3 (red), and CD8 (green). Bar=50μm (low mag), 12.5μm (high mag). (B) Quantification of CD8 T cells (CD3+ CD8+) in the tumors from H- and H+ mice (n=5 each) Unpaired t-test (B) was used for statistical significance (* p<0.05).

**Supplementary Figure 4.**
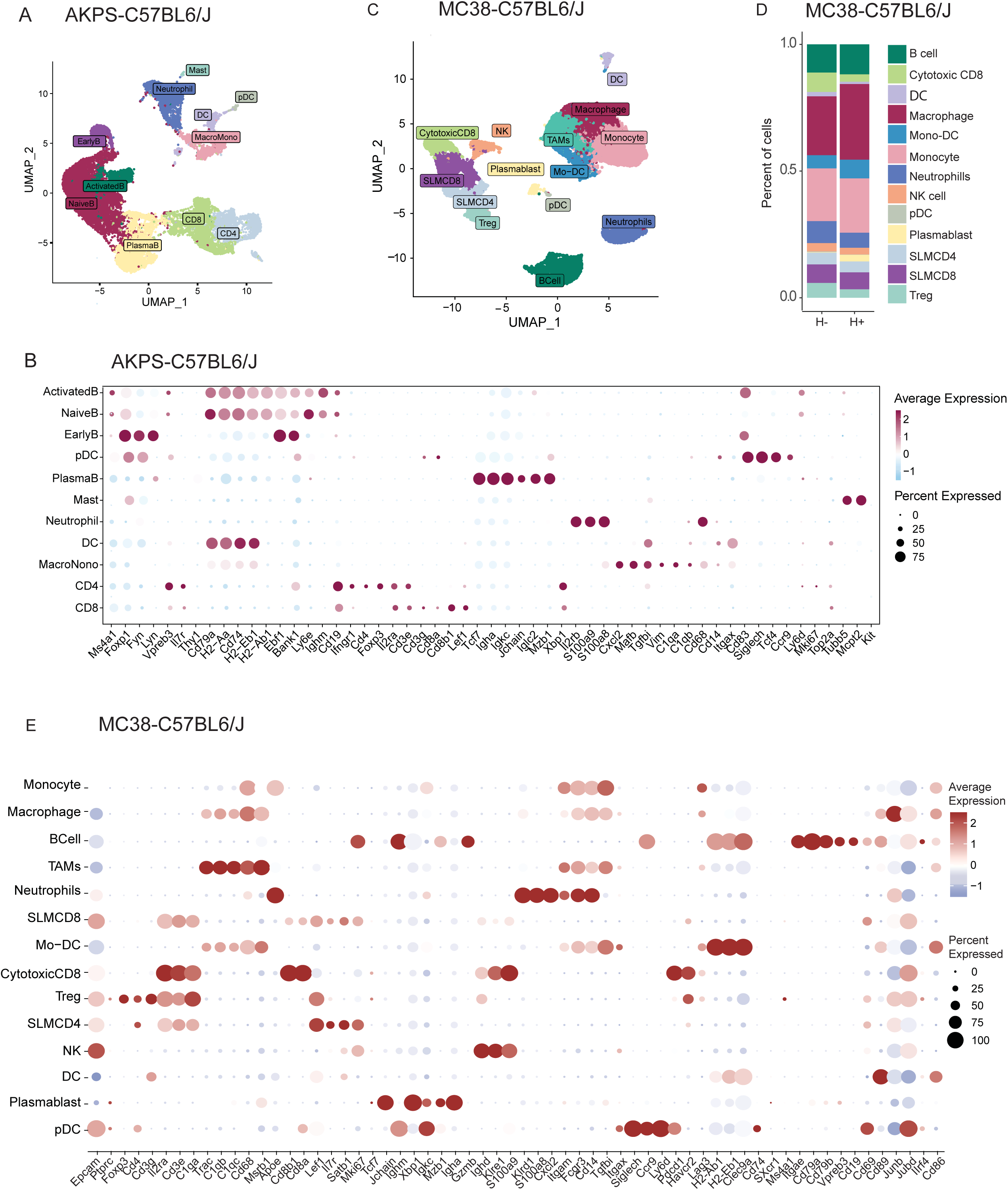
Immune cell cluster classification in AKPS and MC38 tumors. (A) UMAP visualization and clustering of CD45+ cells from digested AKPS tumors (n=4 each). (B) Immune cell cluster classification of CD45+ cells from digested AKPS tumors (n=4 each). (C) UMAP visualization and clustering of CD45+ cells from digested MC38 tumors (n=2 for both H- and H+). (D) Proportions of clustered CD45+ cells of (D). (E) Immune cell cluster classification of CD45+ cells from digested AKPS tumors (n=2 each).

**Supplementary Figure 5.**
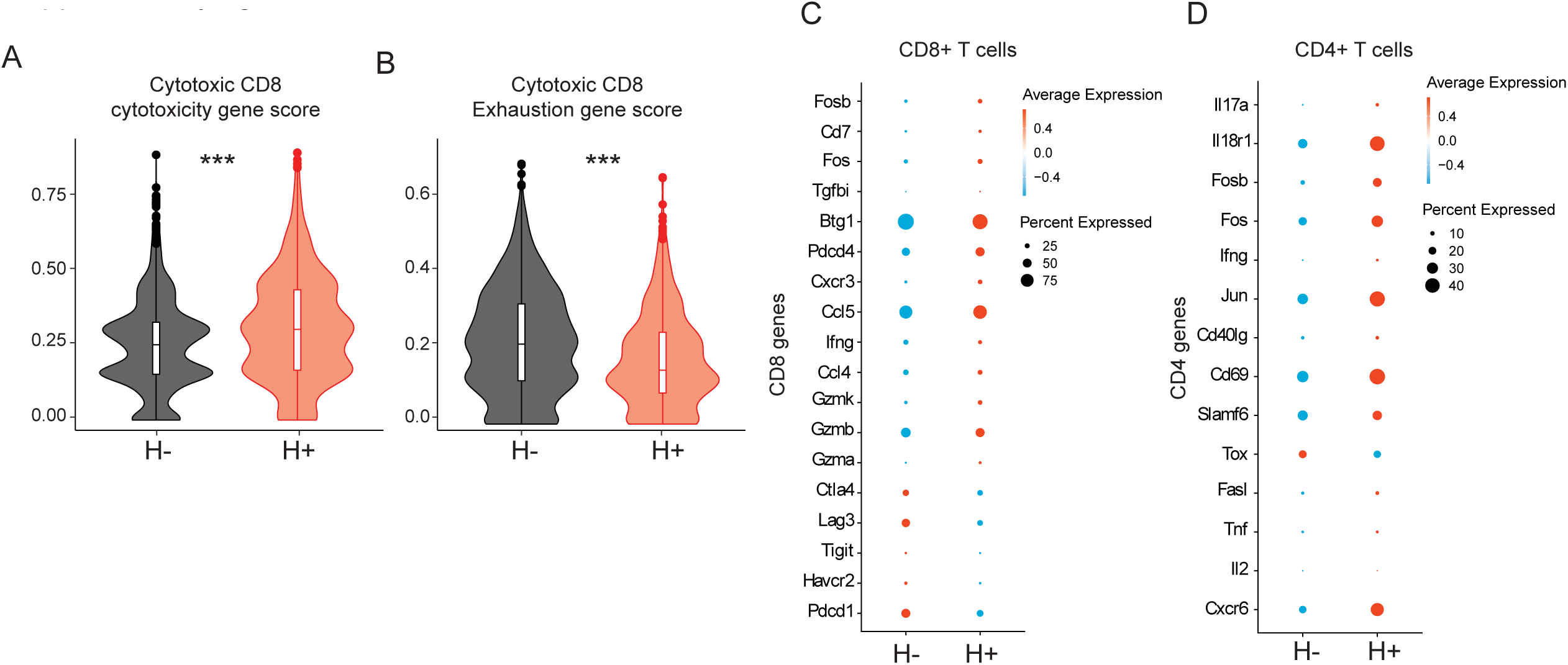
Helicobacter colonization enhances effector gene expression in tumor infiltrating T cells in MC38 tumors. (A) Violin plots of effector scores of the CD8+ T cell cluster. (B) Violin plots of dysfunctional scores of the CD8+ T cells cluster. (C) Bubble plots of immune cell markers in the CD8+ T cell cluster. (D) Bubble plots of immune cell markers in the CD4+ T cell cluster. Unpaired t-test (A-B) was used for statistical significance. For scoring, refer to the method for the list of genes used to score chemotaxis and effector for immune cells.

**Supplementary Figure 6.**
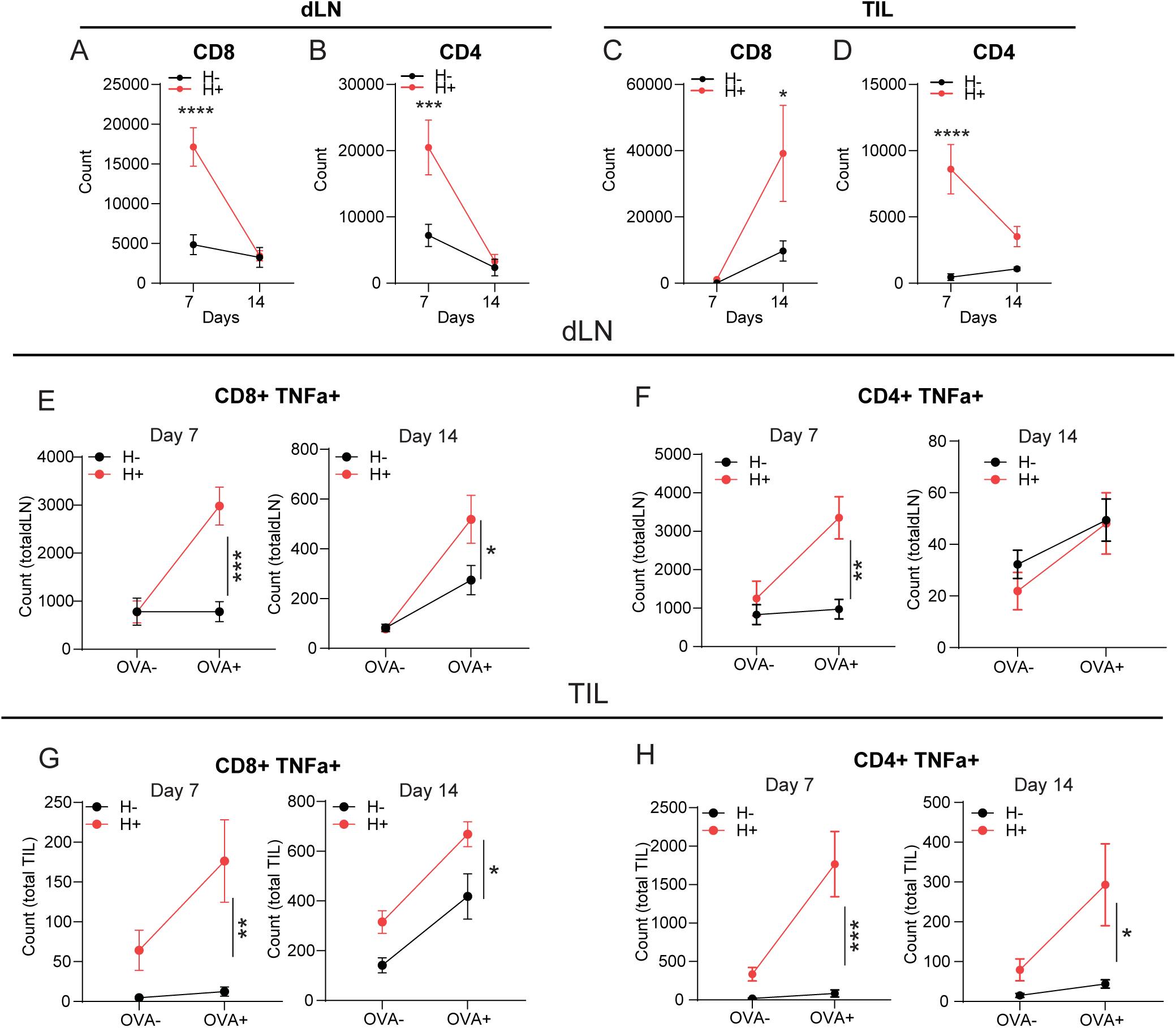
Helicobacter colonization increases CD4^+^ and CD8^+^ T cell abundance following AKPS tumor implantation. (A-D) Quantification of total CD8^+^ and CD4+T cells in draining lymph nodes (dLN) and tumors from AKPS tumor-bearing H- and H+ mice on day 7 (n=5 per group) and 14 (H-, n=8; H+, n=9) post tumor implantation. (E-H) Quantification of tumor-specific (SINFEKL+) CD8+ T cells and CD4^+^ T cells expressing TNFα in dLN (E-F) and tumors (G-H) from AKPS tumor-bearing H- and H+ mice on day 7 (n=5 per group) and day 14 (H-, n=8; H+, n=9), with or without OVA antigen (SIINFEKL) stimulation. Unpaired t-test was used to assess statistical significance. (* p<0.05, ** p<0.01, *** p<0.001, **** p<0.0001).

**Supplementary Figure 7.**
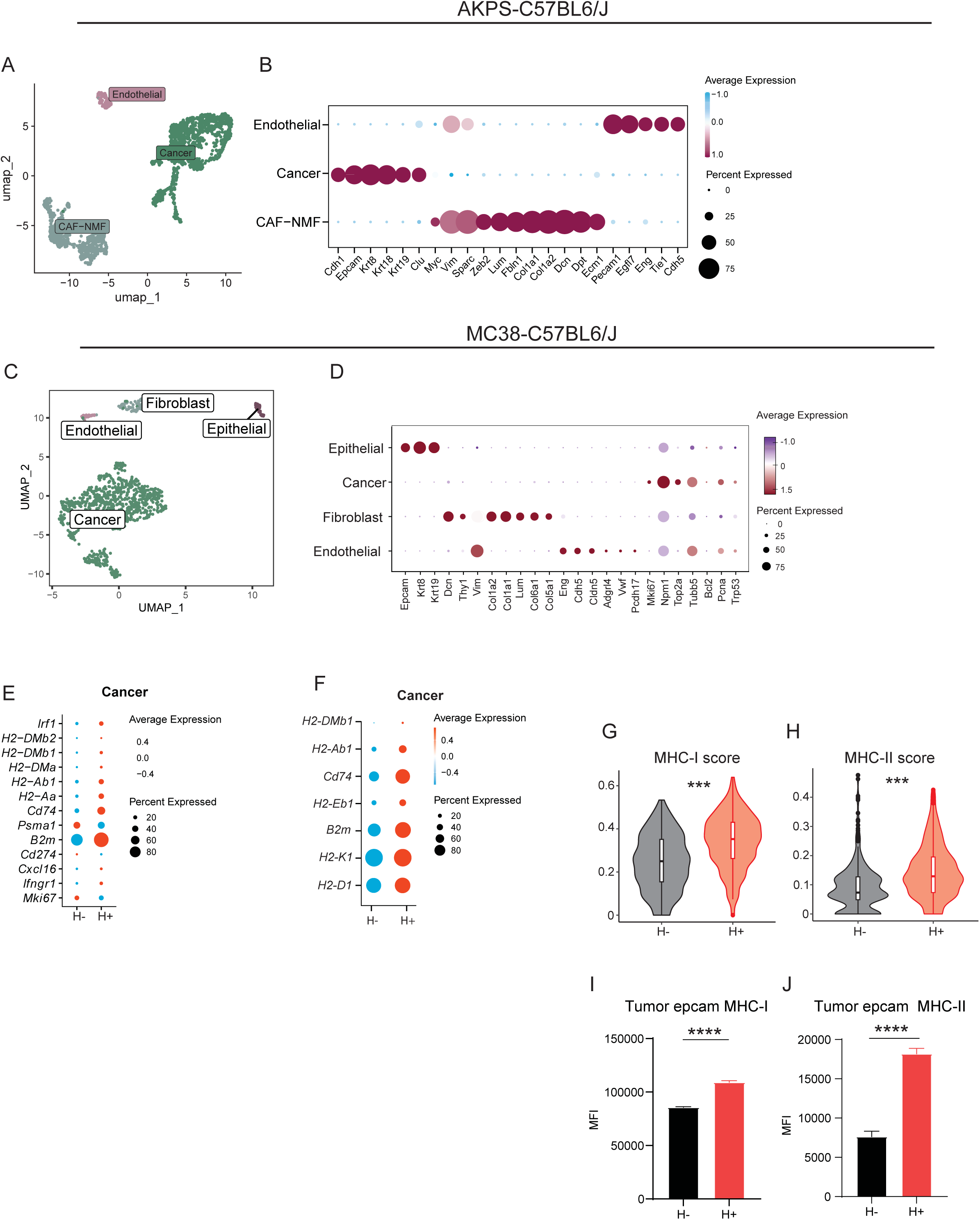
Non-Immune cell cluster classification of AKPS and MC38 tumors. (A) UMAP visualization and clustering of CD45^-^ cells isolated from digested AKPS tumors from H- and H+ mice (n=4 per group). (B) Non-Immune cell cluster annotation of CD45^-^ cells isolated from digested AKPS tumors (n=4 per group). (C) UMAP visualization and clustering of CD45^-^ cells isolated from digested MC38 tumors from H- and H+ mice (n=2 per group). (D) Non-Immune cell cluster annotation of CD45^-^ cells from digested MC38 tumors (n=2 per group). (E) Bubble plots of MHC-related genes in the AKPS cancer cell cluster shown in (B). (F) Bubble plots of cancer-related and MHC-related genes in the MC38 cancer cell cluster shown in (D). (G-H) Violin plots of MHC-I (G) and MHC-II (H) gene score in cancer cell subclusters from MC38 (F; n=2 per group). (I-J) MHC-I (I) and MHC-II (J) expression levels of EpCAM^+^ cells from digested MC38 tumors measured by flow cytometry (H-; n=4, H+; n=8) Unpaired t-test (G-J) was used to assess statistical significance. (* p<0.05, ** p<0.01, *** p<0.001, **** p<0.0001).

**Supplementary Figure 8.**
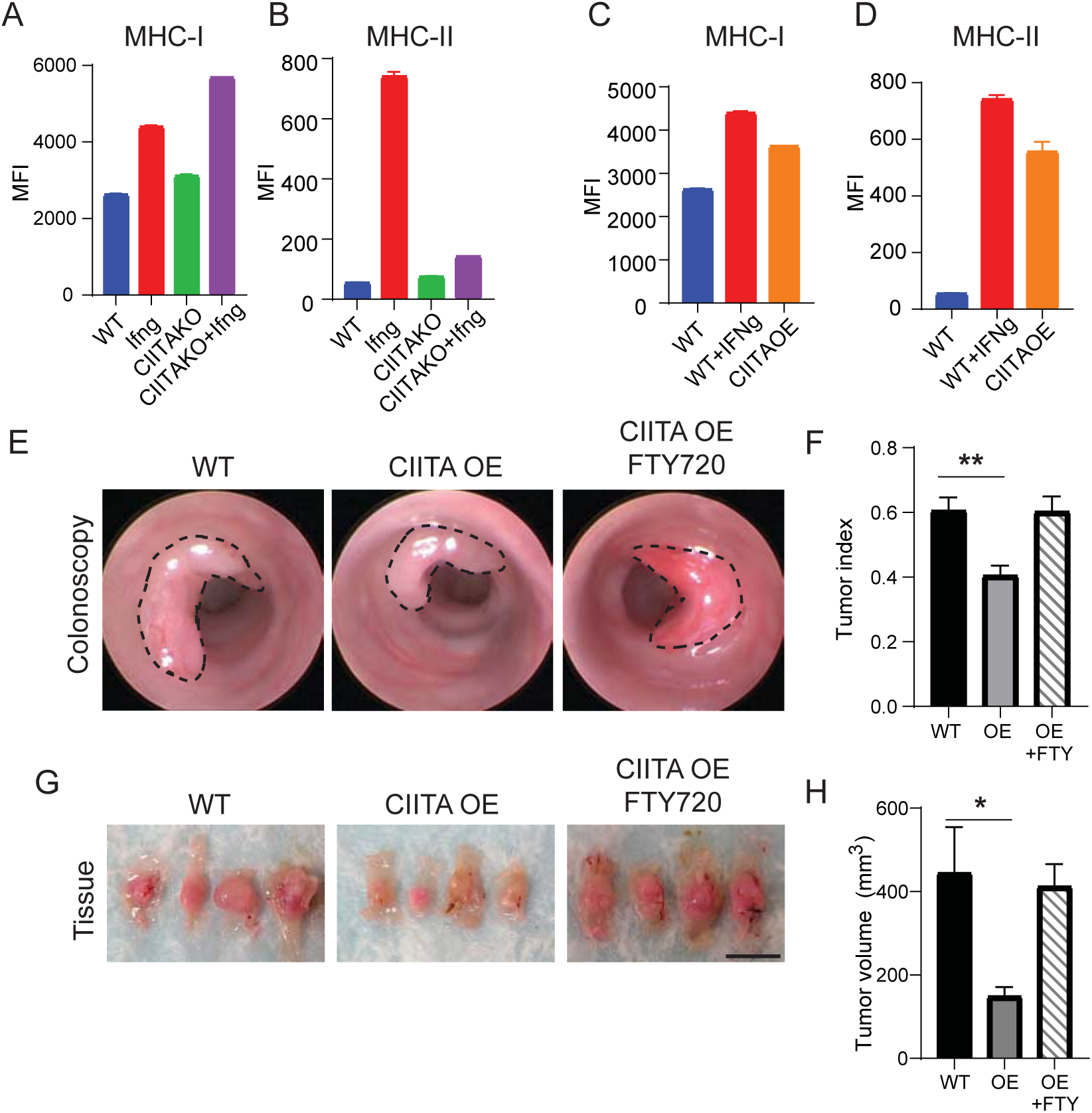
Cancer cell-intrinsic MHC-II requires lymph node-derived immune cells to elicit anti-tumor effects. (A-B) Flow cytometric analysis of MHC-I (A) and MHC-II (B) protein expression in AKPS WT and AKPS CIITA KO with or without IFNγ (2ng/ml for 48 h) (n=3 per group). (C-D) Flow cytometric analysis of MHC-I (C) and MHC-II (D) protein expression in AKPS WT and AKPS CIITA OE at steady state (n=3 per group). (E) Representative colonoscopy images of colon tumors induced by AKPS WT and CIITA OE organoids in H-mice on day 14, with or without FTY720 (Fingolimod; 1 mg/kg, daily oral gavage from day −1 to day 14) treatment. (F) Tumor index of tumors induced by AKPS WT or AKPS CIITA OE organoids with or without FTY720 treatment in H-mice at day 14 (WT, n=5; CIITA OE, n=5; CIITA OE FTY720, n=10). (G) Representative images of gross tumors at day 15 from indicated treatment groups. Scale bar, 1 cm (n=4 per group.) (H) Tumor size measurement corresponding to (E) (WT, n=5; CIITA OE, n=5; CIITA OE FTY720, n=10). Ordinary ANOVA was used for statistical analyses (D, F). (* p<0.05, ** p<0.01).

**Supplementary Figure 9.**
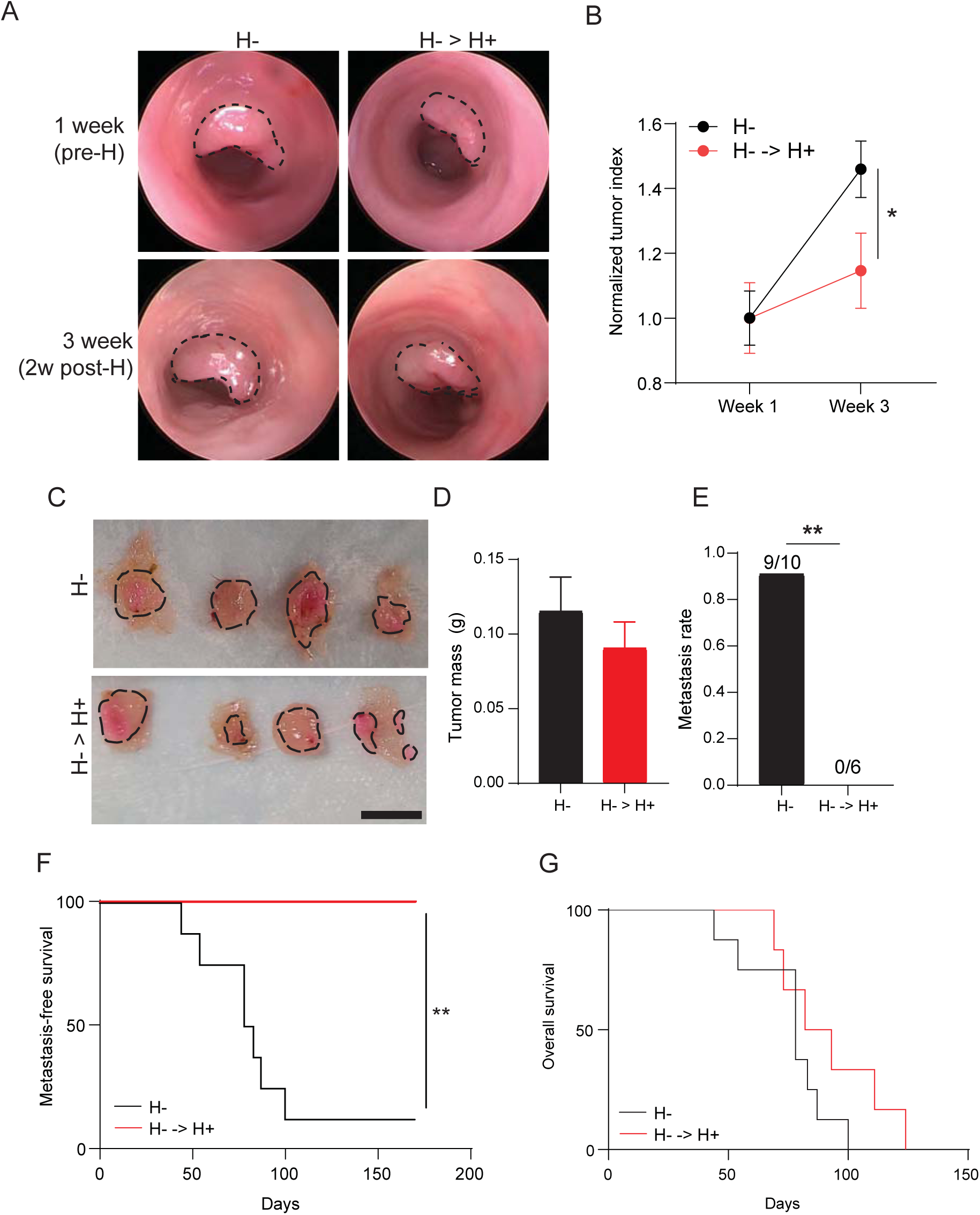
Post-tumor inoculation of *Helicobacter* species confers anti-tumor effects. (A) Representative colonoscopy images of colon tumors induced by AKPS WT organoids in H-or post-tumor colonized (H-→H+) mice at days 7 and 21. Post-tumor Helicobacter inoculation was performed on day 7. (B) Normalized tumor index of AKPS WT induced tumor in H- or H-→> H+ mice at days 7 and 21 (H-, n=4; H---> H+, n=5). Index was normalized to tumor index measurement at day 7 for respective groups. (C) Representative gross images of indicated tumors at day 21 from H- and H-→H+ mice. Scale bar, 1cm. (n=4 per group.). (D) Tumor mass quantification corresponding to (C) (H-, n=4; H--> H+, n=5). (E) Liver metastasis rate in H- and H-→> H+ mice bearing AKPS WT induced tumors (H-, n=10; H-→ H+, n=5). (F) Metastasis-free Survival of H- and H-→ H+ mice bearing AKPS WT induced tumors (H-, n=10; H-→ H+, n= 5). Kaplan-Meier analysis was used for survival comparisons (** p<0.01). (G) Overall survival rate of H- and H-→ H+ mice bearing AKPS WT induced tumors (H-, n=10; H-→ H+, n= 5). Unpaired *t*-test was used for tumor index comparisons (B, D), and Fisher’s exact test were used for metastasis frequency (E). (* p<0.05, ** p<0.01).

**Supplementary Figure 10.**
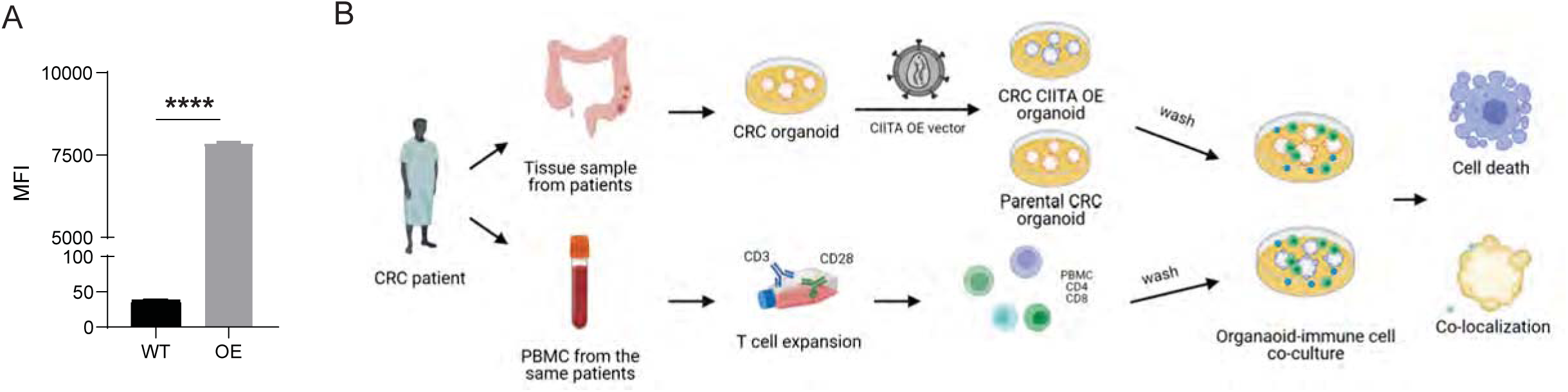
Cancer cell-intrinsic MHC-II enhances cancer-immune interactions and immune-mediated apoptosis in human MSS CRC PDOs. (A) Flow cytometric quantification of MHC-II protein expression in parental (WT) and CIITA overexpression (CIITA OE) human MSS CRC organoids. (n=3 per group). (B) A schematic depicting the human CRC organoid and autologous PBMC co-culture system. Unpaired *t*-test was used for tumor index comparisons (A) was used for statistical significance. (**** p<0.0001).

